# DNA damage checkpoints balance a tradeoff between diploid- and polyploid-derived arrest failures

**DOI:** 10.1101/2025.02.14.638318

**Authors:** Kotaro Fujimaki, Ashwini Jambhekar, Galit Lahav

## Abstract

The DNA damage checkpoint system ensures genomic integrity by preventing the division of damaged cells. This system operates primarily through the G1/S and G2/M checkpoints, which are susceptible to failure; how these checkpoints coordinate quantitatively to ensure optimal cellular outcomes remains unclear. In this study, we exposed non-cancerous human cells to exogenous DNA damage and used single-cell imaging to monitor spontaneous arrest failure. We discovered that cells fail to arrest in two major paths, resulting in two types with distinct characteristics, including ploidy, nuclear morphology, and micronuclei composition. Computational simulations and experiments revealed strengthening one checkpoint reduced one mode of arrest failure but increased the other, leading to a critical tradeoff for optimizing total arrest failure rates. Our findings suggest optimal checkpoint strengths for minimizing total error are inherently suboptimal for any single failure type, elucidating the systemic cause of genomic instability and tetraploid-like cells in response to DNA damage

**Highlights:**

- Arrest-failed cells result from two routes, each leading to a distinct type
- These types differ in nuclear morphology, size, ploidy and micronuclei composition
- Strengthening one checkpoint reduces one arrest-failure type but increases the other
- Total error is minimized when checkpoint strengths are suboptimal for each type

**eTOC:** Single-cell quantitative analyses and computational simulations reveal that the DNA damage checkpoint system balances a tradeoff between reducing arrest failure from diploid cells and from damage-induced polyploid cells. The study suggests that to minimize total arrest failure, the checkpoint strengths need to be sub-optimal for each individual arrest-failure type.

## INTRODUCTION

Failure analysis is a process often used in engineering to determine how a specific system underperforms. Identifying the fundamental causes of failure can guide corrective actions to improve performance quality and provide insights into the system’s working mechanisms. Similar to engineering, various cellular functions can fail in cells, leading to a range of pathophysiological conditions. Thus, a systematic analysis of how a certain cellular function underperforms in healthy cells can elucidate the underlying cause of such malfunction as well as its mechanism of action. Here we analyzed failures of the DNA damage checkpoints to better understand their function and quantitative constraints.

The DNA damage checkpoint system prevents cells from replicating their DNA and dividing in the presence of DNA damage^1^. Such a system is critical for cell viability, as cells routinely experience DNA damage from both endogenous and exogenous sources, such as DNA replication error, genotoxic chemicals, or radiation. Checkpoint failure can lead to catastrophic genome instability and pathophysiological conditions such as cancer^2^. The system functions primarily through two checkpoints that occur at the G1 and G2 phases of the cell cycle^1,3^. The G1/S checkpoint prevents cells from progressing to S phase by driving cell cycle exit into G0^4,5^. This prevents chromosome breaks^6^ and blocks DNA damage-induced whole genome duplication^7–9^. The G2/M checkpoint halts damaged G2 cells from undergoing aberrant mitoses^10^ which can result in micronuclei^11^ – small nuclear compartments separate from primary nuclei containing chromosome fragments or whole chromosomes^12^. Prolonged G2/M checkpoint arrest can also cause G2 arrested cells slip into a G0 or G1 state without mitosis^8,13^, a phenomenon known as mitotic skipping^14,15^.

The molecular mechanisms underlying these checkpoints have been extensively studied and involve both post-transcriptional and transcriptional regulation. The G1-S checkpoint is initiated by the damage-sensing kinase ATM, which phosphorylates numerous targets that suppress cell-cycle progression^16,17^. The G2-M checkpoint relies primarily on the kinase ATR^10,16^, although ATM can also contribute to its initial activation^10,17,18^. The two checkpoints contain shared downstream targets, with the tumor suppressor p53 being a key component operating in both pathways, enforcing longer-term arrest by transcriptional activation of its targets including cyclin-dependent kinase inhibitor proteins^18,19^.

The two checkpoints exhibit distinct vulnerabilities in terms of sensitivity, speed, and robustness. The G1/S checkpoint is sensitive to low levels of DNA damage, responding to as little as a single double-stranded break^20^. However, it is slow in commitment (2-4 hours) ^6,21–24^ due to the time required to induce sufficient cyclin-dependent kinase inhibitor proteins such as p21 to overcome persistent cyclin-dependent kinase activity^5^. The G1/S checkpoint is also leaky, resulting in escape from arrest^6,25^, due to stochastic reduction in p53 levels and the concomitant drop in p21^25^. In contrast, the G2/M checkpoint is rapid in commitment (less than 0.5 hour) ^21^ but insensitive to low levels of DNA damage, requiring 10 to 20 double-stranded breaks to become activated^26,27^. It has been proposed that these distinct vulnerabilities in the checkpoints contribute to arrest failures and genome instability^22,28^.

Many studies of DNA damage checkpoints have evaluated each checkpoint individually and have monitored cells for relatively short periods of time (hours). However, a quantitative understanding of how the two DNA damage checkpoints operate together to guide and optimize cellular outcomes is still missing. Inspired by failure analysis, we exposed non-cancerous human cells with intact checkpoints to exogenous DNA damage and monitored spontaneous arrest failure over several days. We identified two distinct modes of arrest failure, each caused by a different route. Using a combination of single cell imaging and computational modeling, we demonstrate that strengthening one checkpoint reduced the occurrence of one mode of arrest failure but increased that of the other. Our study suggests that the two checkpoint systems can minimize total error only when the checkpoint strengths are suboptimal for each arrest-failure type.

## RESULTS

### Checkpoints failures are highest under intermediate levels of DNA damage

To quantify arrest failure in cells with functional checkpoints, we exposed an immortalized non-cancerous human retinal epithelial cell line (hTERT RPE-1, hereafter referred to as RPE) to various sub-lethal doses of irradiation or DNA damaging drugs and examined micronucleation, a marker of cells that have undergone division with unresolved DNA damage^12^ (Fig. 1A, Fig. S1A). Micronuclei were detected using a newly developed pipeline that differentiates primary nuclei (PN) from micronuclei (MN) through machine-learning-based pixel categorization (Fig. 1B, see *Methods* for detail). The relationship between the dose of insult (irradiation or drug) and the amount of DNA damage (measured by γ-H2AX foci) was monotonic, with higher doses leading to more damage (Fig. S1B). Micronucleated cells were detected across all tested conditions, but their relationship with DNA damage dose was not linear. Specifically, the fraction of micronucleated cells was maximal under intermediate levels of DNA damage, resulting in a bell-shaped dose-response curve (Fig. 1C). The mean number of treatment-induced γ-H2AX foci at which the maximal fraction of micronucleation occurred was similar across different DNA-damaging agents (Fig. S1C), ruling out reagent-specific off-target effects. These findings suggest the existence of a DNA damage “Goldilocks Zone” that elicits checkpoint failure and hence micronucleation in normal human cells.

**Figure 1:**
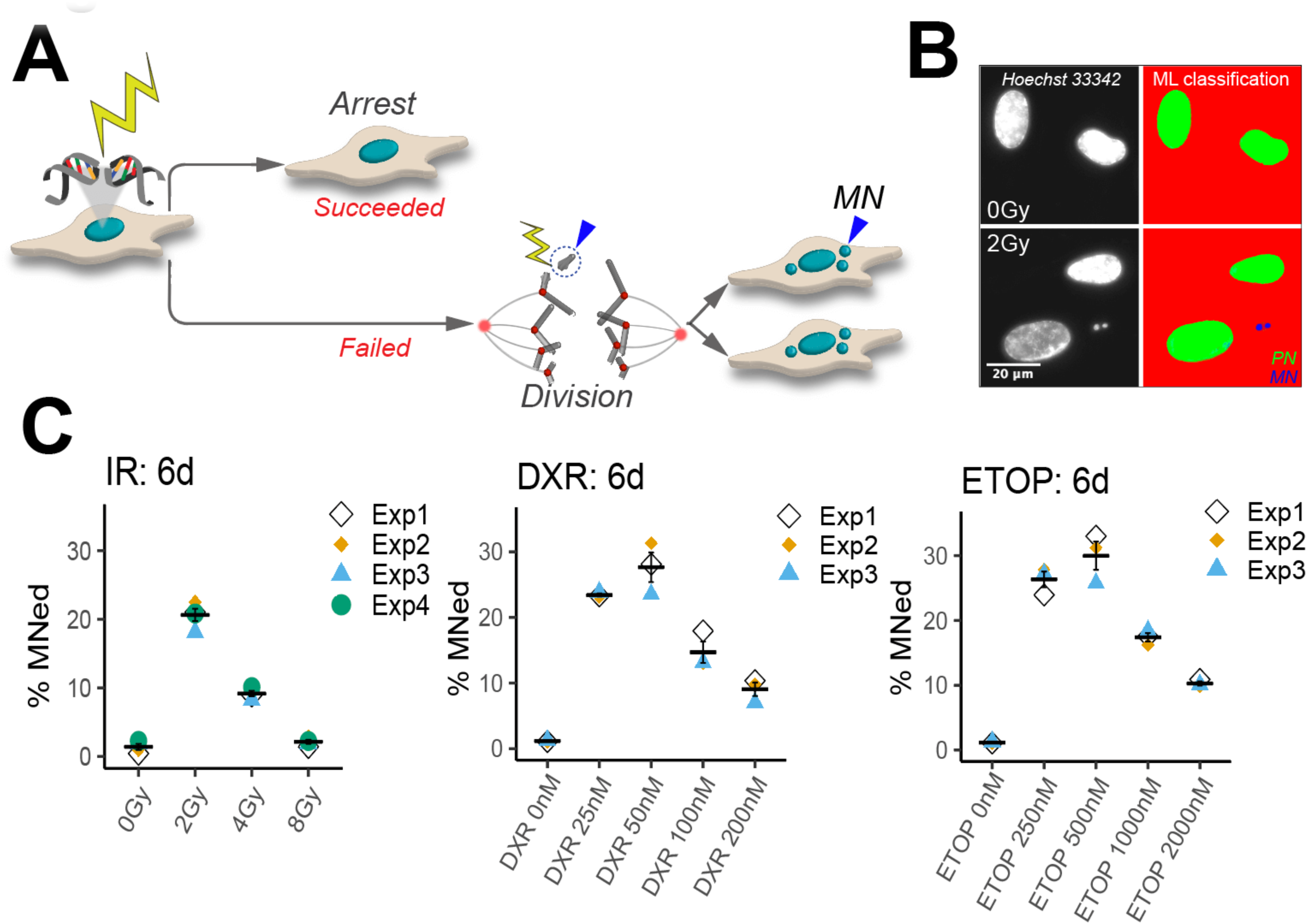
The fraction of micro-nucleated cells is highest under intermediate levels of DNA damage. **(A)** Schematic of the cellular outcomes following DNA damage. Cells with functional checkpoints arrest in the cell cycle to prevent division with unresolved DNA damage. When arrest fails, subsequent division with unrepaired DNA damage could result in micronucleation. MN, micronuclei. **(B)** Left – images of DNA stained RPE1 cells. Right - quantification of the primary nuclei (PM, green) and micronuclei (MN, blue) using machine-learning-assisted nuclear segmentation. **(C)** Percentage of micronucleated cells across different doses of various DNA-damaging agents (n ≥ 3 biological replicates with >250 cells each). Irradiated cells were cultured for 6 days. Cells were exposed to one of the topoisomerase inhibitors for 2 days then recovered for 4 days before imaging. IR, irradiation; DXR, Doxorubicin; ETOP, Etoposide. MNed, micronucleated. Error bar, SEM. Shape symbols indicate independent biological replicates.

### Nuclear morphology elucidates two distinct types of arrest-failed micronucleated cells

A subset of micronucleated cells under DNA-damaged conditions showed notable deformation in their primary nuclei, deviating from the typical oval shape seen in non-damaged RPE cells (Fig. 2A). We termed the degree of nuclear deformity “irregularity” and calculated an irregularity score for each cell based on the divergence of its area from a convex area (Fig. 2B). We noticed that the irregularity score, as well as other nuclear features such as size, varied remarkably among micronucleated cells (Fig. 2C, Fig S2A, B). To identify potential correlation between various nuclear features, we performed principal component analysis (PCA) and clustering analysis on several nuclear features – including irregularity score, number of micronuclei, number of γ-H2AX foci, nuclear area, and DNA content. This analysis revealed two clusters within the micronucleated subset (Fig. 2D, Fig. S2C, D). Cells in cluster 1 (hereafter Type 1 cells) were characterized by low irregularity scores, relatively few micronuclei, and DNA content that was mostly 2C, whereas cells in cluster 2 (hereafter Type 2 cells) were characterized by high irregularity scores, high levels of micronucleation, large primary nuclei, and mostly 4C DNA content (Fig. 2E, Fig. S2E, F). The number of γ-H2AX foci did not vary significantly between the two types (Fig. 2E, Fig. S2E-G). These results suggest that arrest-failed micronucleated cells fall into two major categories (Fig 2F), and that the levels of DNA damage do not correlate with these two phenotypes.

**Figure 2:**
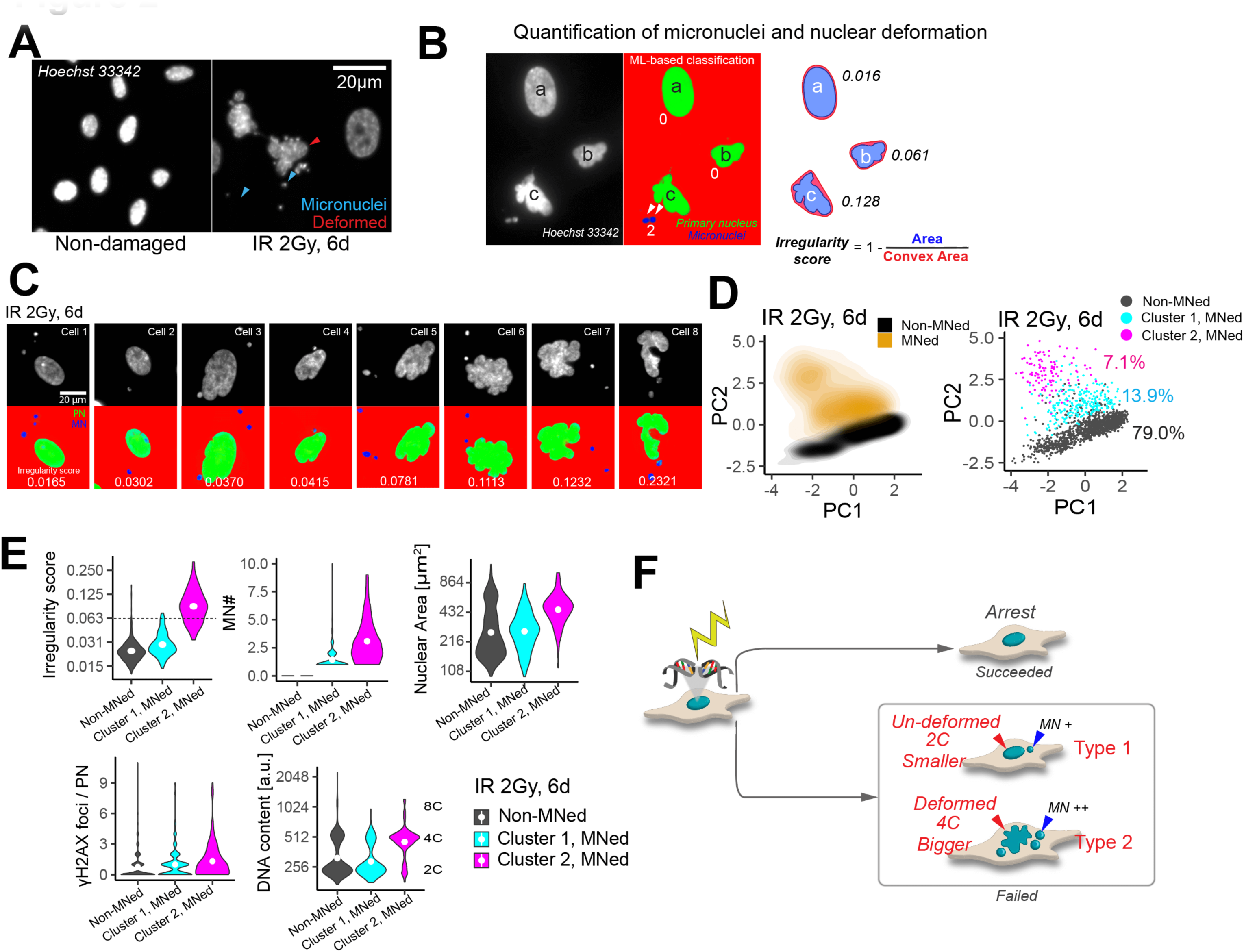
Nuclear morphology elucidates two distinct types of arrest-failed micronucleated cells. **(A)** Nuclear morphologies of RPE cells 6 days after treated with irradiation. Colored arrowheads indicate deformed primary nuclei (red arrow) and micronuclei (blue arrows). **(B)** Quantification of nuclear deformation with irregularity score (one minus convex area over area). **(C)** Examples of micronucleated cells with varying degrees of nuclear deformity. Cells were treated as in A. **(D)** 2D Scatterplot of PC1 and PC2 of irradiated cells (left) and the two clusters within the micronucleated subset (right) (n > 1600 cells). Irregularity score, micronuclei number, nuclear area, γ-H2AX foci, and DNA content were quantified for each nucleus. Principal component analysis (PCA) was performed on these five features. k-means clustering (k=2) was performed on micronucleated cells. **(E)** The distribution of the five nuclear features used for the PCA analysis in **D** in non-micronucleated cells, micronucleated cluster 1 cells, and micronucleated cluster 2 cells (n > 1600 cells) – data from D. Dashed line indicates the top 2 percentile of the irregularity scores in non-damaged conditions (top left). White dot indicates mean. **(F)** Schematic summary of the main features differing between the two nuclear phenotypes observed among arrest-failed micronucleated cells following DNA damage.

### Arrest-failure phenotypes appear immediately after division but are predetermined prior to division

We next sought to identify the cellular events preceding the formation of Type 1 and Type 2 cells. Since irradiation, Doxorubicin, and Etoposide produced a similar spectrum of nuclear morphologies with similar dose-response curves, we used irradiation as a representative DNA damaging agent for further studies. Cells expressing an NLS-CFP nuclear marker were treated with 2Gy of irradiation (a dose that yielded maximal frequency of deformed nuclei) and the extent of nuclear deformity (higher for Type 2 cells) was monitored using live-cell imaging. Micronucleated cells appeared after mitosis, concurrently with a subset of them exhibiting deformed nuclei (Fig. 3A-B), indicating that both arrest-failure phenotypes appear immediately after division. Irregularity scores varied dramatically between cells (Fig. 3B-C), but remained relatively stable for each individual cell throughout interphase. Irregularity scores of sister cells were strongly correlated (Fig. 3D), suggesting that the susceptibility to nuclear deformation arose in the mother cell prior to its division. Pre-arresting cells with CDK4/6 inhibitor Palbociclib or MDM2 antagonist Nutlin-3 before irradiation (Fig. S3A, C) eliminated nuclear deformation and micronucleation following damage (Fig. S3B, D), indicating causality between division and these features. Together, these results suggest that Type 1 and 2 cells are generated upon divisions, and these arrest-failure phenotypes are predetermined prior to division.

**Figure 3:**
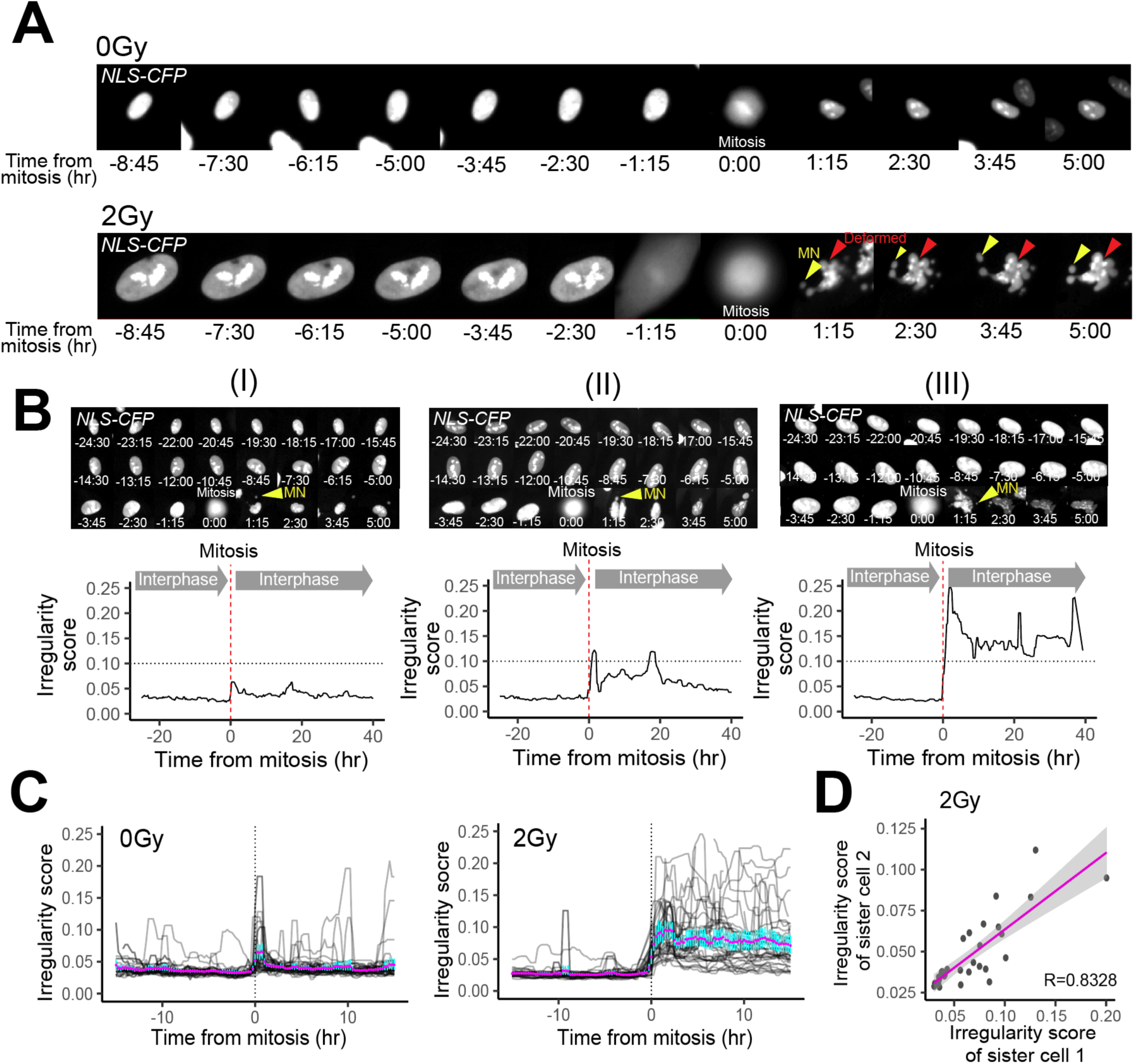
Arrest-failure phenotypes appear immediately after division but are predetermined prior to division. **(A)** Time-lapsed images of nuclear morphology before and after mitosis in a non-damaged control (top) or an irradiated (bottom) cell expressing nuclear-localized CFP. MN, micronuclei. **(B)** Time-lapsed images of nuclear morphology before and after mitosis in three examples of irradiated cells. Yellow arrowhead indicates micronuclei. The bottom panel shows real-time quantification of irregularity scores in the corresponding cells. Time of mitosis is indicated by a red dotted line. Fluctuations arise from overlap of adjacent cells or nuclear movement. The horizontal dashed line intersecting at 0.1 was plotted to ease the comparison of irregularity scores between the three examples. **(C)** Quantification of irregularity scores over time. 40 cells were randomly sampled for each condition. Cells were synchronized *in silico* according to their mitotic timing. The mean and its confidence interval calculated from bootstrapping are respectively shown in magenta dots and cyan lines. **(D)** Irregularity scores of sister cells, quantified from live imaging of NLS-CFP expressing cells (n = 70 daughters).

### Ploidy prior to division determines the arrest-failure phenotype

DNA damage or stress can trigger polyploidization in normal human cells^29–32^ (Fig. 4A, Fig. S4A). Since Type 2 cells exhibit higher DNA content and larger nuclear size (Fig. 2E), we hypothesized that these cells result from division of damage-induced polyploid cells (Fig. 4B). To investigate this hypothesis, we monitored divisions in live cells expressing an S/G2 reporter hGeminin-mCherry and a nuclear marker, NLS-iRFP, and assessed their DNA content at the endpoint (Fig. 4C). Newborn daughters were identified as cells that (i) resulted from a division; and (ii) remained negative for Geminin after division (i.e., in G0 or G1) (hereafter Divided & Geminin-cells). Without DNA damage, cells had either 2C or 4C DNA content, while all Divided & Geminin-cells had 2C DNA content (Fig. 4D). Notably, following irradiation, the Divided & Geminin-population contained cells with both 2C and 4C DNA content (Fig. 4D), suggesting that the latter subset of cells originated from mother cells with 8C DNA content. 4C daughters were associated with higher irregularity scores (Fig. 4E) and with larger nuclear size (Fig. S4B), both characteristics of Type 2 cells. Furthermore, compared to cells with undeformed nuclei (enriched for Type 1), cells with deformed nuclei (enriched for Type 2) exhibited higher percentage of centromeres in micronuclei (Fig. 4F, G), consistent with the known propensity of polyploid division for chromosome missegregation^33–35^ (Fig. S4C). Taken together, our data suggest that the two main phenotypes resulting from checkpoint failures post DNA damage depend on the ploidy of the dividing mother cell: cells with non-deformed smaller nuclei (Type 1) result from division of diploid cells, while cells with deformed larger nuclei (Type 2) result from division of polyploid cells (Fig. 4H).

**Figure 4:**
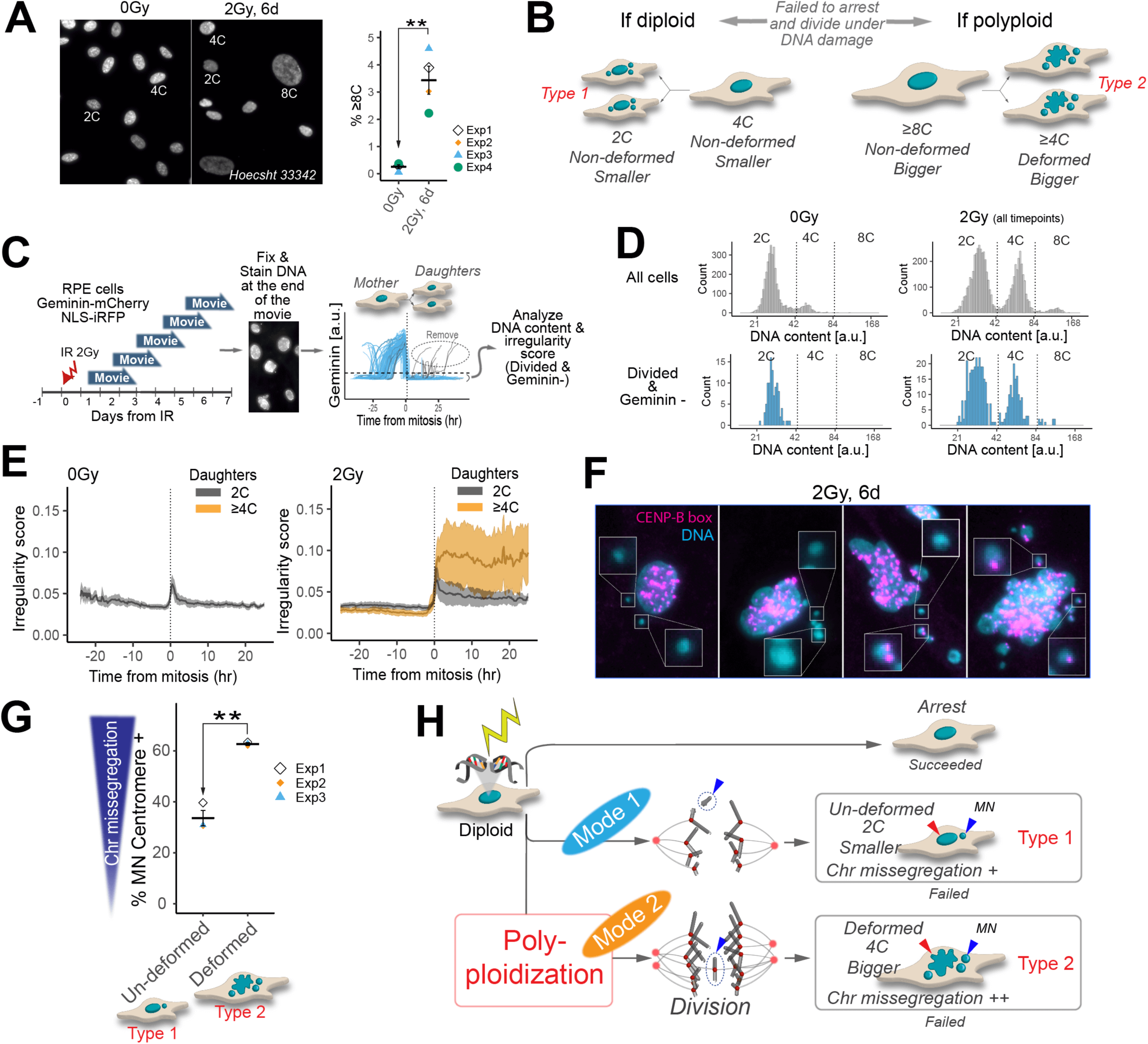
Ploidy prior to division determines the arrest-failure phenotype. **(A)** Left: nuclear images in non-damaged and damaged conditions, including a polyploid nucleus in the latter. Right: percentage of ≥8C polyploid cells (n = 4 with >630 cells per condition) – data from Fig. 1C. **P < 0.01, 1-taild paired t test. Error bar, SEM. **(B)** Model for the impact of mother cells’ ploidy on nuclear deformation and micronucleation post-division in the presence of DNA damage. **(C)** Schematic of the live-cell imaging in combination with DNA content analysis. To evenly capture divisions following irradiation, cells with Geminin and NLS reporters were imaged in different time windows following irradiation. Non-irradiated cells were imaged as a control. At the end of each movie, the cells were fixed and stained for their DNA. Irregularity scores and the endpoint DNA content were analyzed for cells that divided and remained negative for Geminin (referred to as Divided & Geminin-cells.) **(D)** Quantification of DNA content in all cells (top) and in Divided & Geminin-cells (bottom) under undamaged or irradiated conditions (n > 3500 cells per condition). **(E)** Real-time quantification of irregularity scores in 4C and 8C mothers and their daughters (n = 82, 237, and 137 cells correspondingly for non-damaged 4C, irradiated 4C, and irradiated 8C subsets). The cells were synchronized *in silico* according to their mitotic timing. The median and interquartile range are respectively shown in thick lines and colored ribbons. **(F)** Centromere detection using peptide nucleic acid fluorescence in situ hybridization assay (PNA FISH) for CENP-B box DNA repeats (see *Supplementary method* for details). Micronucleated cells with varying degrees of nuclear deformation are shown. **(G)** Percentage of micronucleated cells with micronuclear CENP-B box signal, in cells exposed to 2Gy of irradiation and cultured for 6 days (n=3 biological replicates with >2300 cells each). Cells were categorized into un-deformed or deformed primary nuclei. Error bar, SEM. **P < 0.01, 1-taild paired t test. **(H)** Schematic of the two modalities in arrest failure in cells with functional checkpoints. Damage-induced polyploidization of arrest-failed cells post-damage affect the choice between Type 2 or Type 1 outcomes.

### Polyploidy giving rise to Type 2 cells originates from G0 cells with 4C DNA content

Damage-induced polyploidy is known to result from endoreplication^7–9^ of G2 cells that entered a 4C G0/G1-like state without intervening mitosis^14,15^ (i.e., mitosis skip). However, whether cells enter a G0 or G1 state with 4C DNA prior to whole genome duplication remains elusive. Thus, we next investigated the cell-cycle phases en route to damage-induced polyploidy resulting in Type 2 cells. We inferred cell-cycle phase transitions by immunostaining endogenous Cdt1 and Geminin and quantifying pulse-labeled thymidine analog EdU. The simultaneous labeling of these three markers can differentiate between all the non-mitotic phases (G0, G1, S, and G2) (Fig. 5A, S5A). We agnostically extracted >100 features from immunostaining images taken at different time points after irradiation (Fig. 5B) and visualized changes in cellular profiles by dimensionality reduction (Fig. 5C, see *Methods* for detail). In the absence of irradiation, cells exhibited a G1, S, or G2 signature of the three markers (i.e., Cdt1, EdU, and Geminin) with the corresponding 2C, 2C∼4C, and 4C DNA content, as expected (Fig. 5C, (I) 0d). To infer the polyploidization route before division, we inferred the trajectory^36^ in non-micronucleated cells with ≥4C DNA content, using the quantified features. The inferred trajectory aligned closely with the sequence of sampling days (Fig. 5C, (IX) vs. (X)). Trajectory analysis revealed that, within a day after irradiation, cells transitioned from (a) a G2 state; to (b) an unorthodox Cdt1 positive state with low (rather than absent) Geminin; to (c) a 4C G0 state lacking both Cdt1 and Geminin (Fig. 5C, (II): transition from ⓐ, ⓑ, to ⓒ), suggesting a G2-to-G0 transition. Consistently, cells in (c) lacked phosphorylated Rb, which marks cycling cells (Fig. S5B). Polyploid cells were not detectable at this stage (Fig S5C, 1 day). After 2 or more days, some of the 4C G0 cells transitioned to ≥8C polyploid state (Fig. 5C, (IV) 4d and (X)). These results suggest that the polyploid cells giving rise to Type 2 cells with deformed nuclei emerged from G2 cells that transitioned to a 4C G0 state prior to re-entering the cell cycle and DNA replication.

**Figure 5:**
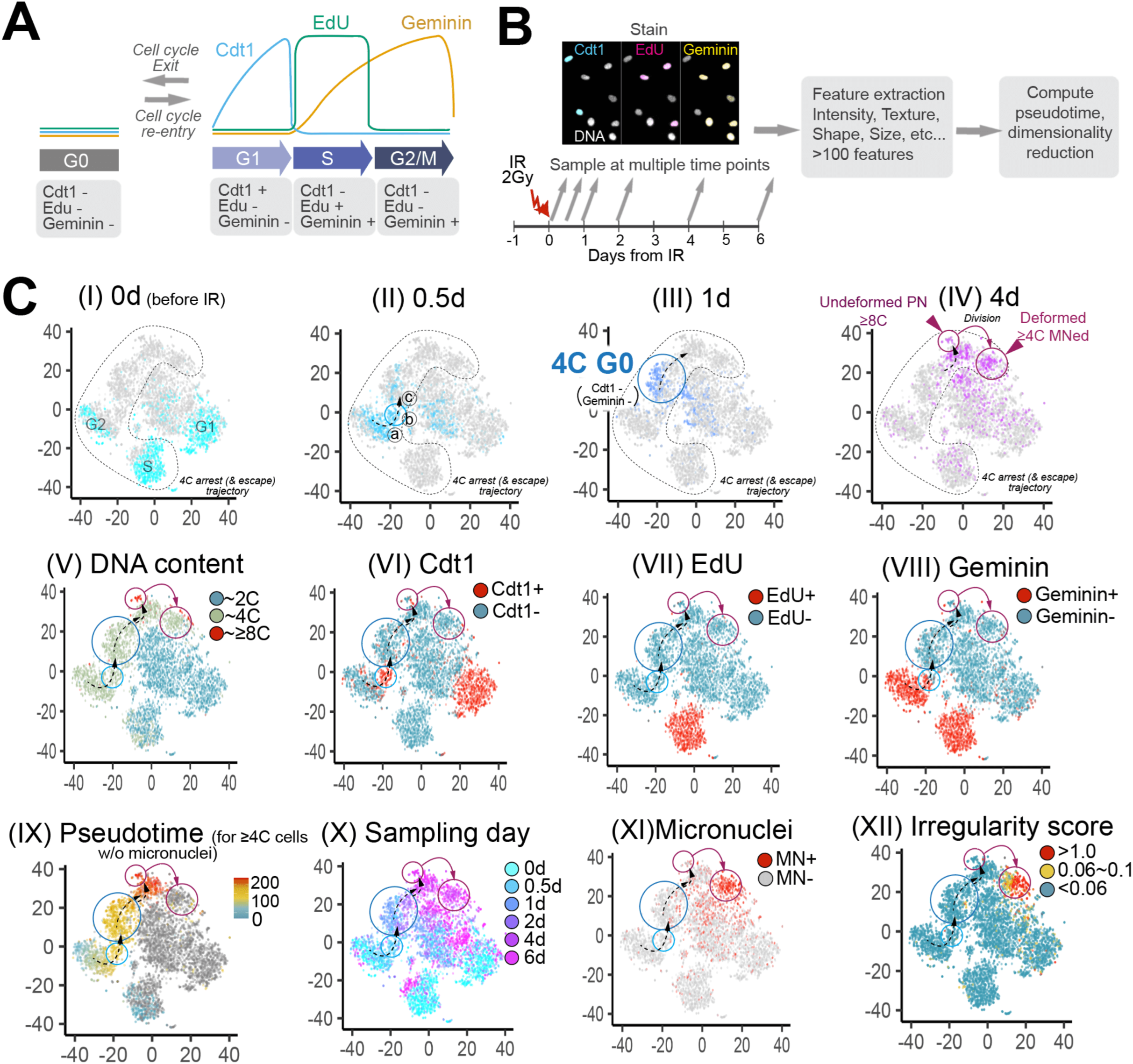
Polyploidy giving rise to Type 2 cells originates from G0 cells with 4C DNA content. **(A)** Schematic showing the expression of endogenous Cdt1 and Geminin and pulse-labeled EdU signal in each cell-cycle phase. Endogenous Cdt1 and Geminin are down-regulated in G0^46^ due to their transcriptional reliance on E2F activators^47^. **(B)** Experimental and analytical scheme for deducing cell-cycle state transitions. Cells were EdU pulse-labeled and stained for endogenous Cdt1 and Geminin. More than 100 features were agnostically extracted from 0- to 6-day irradiated cells for trajectory inference. **(C)** Molecular and morphological profile of 0- to 6-d irradiated cells (n > 570 cells per timepoint). Each dot in the t-SNE plots represents a single cell. On the top row, cells from the indicated post-irradiation day are highlighted in color. Using the extracted features, pseudotime was estimated for ≥4C non-micronucleated cells, from which the 4C arrest (& escape) trajectory was deduced. Dashed arrows and circled solid lines were drawn to clarify the deduced temporal order of events. Cdt1, EdU, and Geminin showed bimodal distributions (Fig. S5A), and thus their “positive” or “negative” expression was annotated for each cell. MN+, cells with micronuclei; MN-, cells without micronuclei.

### Checkpoints stringencies balance a tradeoff between diploid- and polyploid-derived arrest-failure types

To understand how the DNA damage checkpoint system collectively affects the two arrest-failure phenotypes, we created a computational cell-cycle-checkpoint model. We constructed a stochastic Markov-chain-like model, in which cells transition between cell-cycle phases based on probabilities that could be affected by DNA damage checkpoints (Fig. 6A). Higher damage increases the likelihood of checkpoint activation (see *Methods* for detail). The G1/S checkpoint promotes arrest by promoting cell-cycle exit from G1^4,5^ and by preventing the escape from G0 (Fig. 6A). The G2/M checkpoint prevents G2 progression and allows slippage into G0 (hereafter called “cell-cycle reset,” differentiating from the regular G1-to-G0 cell-cycle exit; Fig. 6A). Based on data collected here and by others^33–35^, we determined that division with damage or polyploidy drives mitotic error (represented by DSB- or chromosome missegregation-driven micronucleation), and that diploid and polyploid daughters with mitotic error represents Type 1 and 2 cells (collectively arrest-failed cells), respectively (Fig. 6A). Model parameters were set according to experimental observations (Fig. 6A, Supplementary Table 1).

**Figure 6:**
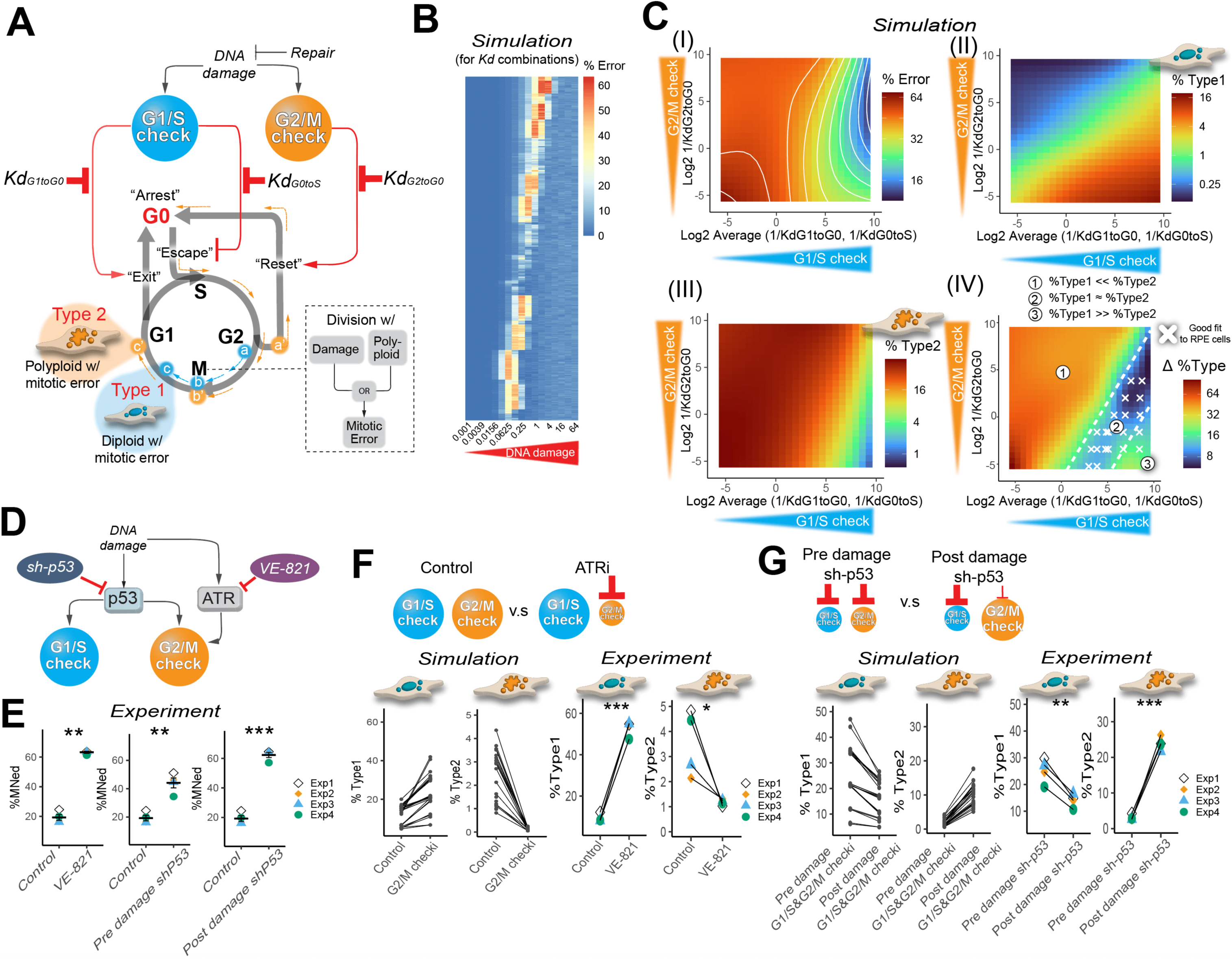
Checkpoint system faces a tradeoff between the diploid- and polyploid-derived arrest-failures. **(A)** A framework of the Markov-chain-like model. We defined that cells escape G0 and reenter the cell cycle via S phase (and not G1 phase), similar to E2F activators driving cell-cycle reentry via committing cells to S phase^65,66^. We assume that division with damage or polyploid state can result in mitotic error (see the main text, Fig. S5A, and *Methods* for details). Diploid and polyploid cells with mitotic error are defined as Type 1 and Type 2 cells, respectively; “Reset-and-Escape” followed by erroneous mitosis (a’ to b’) give rise to polyploid Type 2 cells (c’) whereas erroneous mitosis without “Reset-and-Escape” (e.g., a to b) result in diploid Type 1 cells (c). **(B)** Simulated total percentage of mitotic error under a range of DNA damage levels (columns) for each of the 729 unique *Kd* combinations (rows); *Kd* was varied between 0.001 and 64. Result from the 6^th^ model day after the induction of damage. Percentage of mitotic error was calculated from three sets of stochastic simulations (100 seeding cells each). **(C)** Simulated effect of the G1/S and G2/M checkpoints’ stringency on arrest failure types – simulation from Fig. 6B. Arrest-failure phenotypes were quantified for the intermediate DNA damage level that resulted in the highest percentage of mitotic error (the most vulnerable DNA damage level). Result from the 6^th^ model day after the induction of damage, each with three sets of sets of stochastic simulations with 100 seeding cells each. The stringency of the G1/S checkpoint, the average of 1/*Kd_G1toG0_* and 1/ *Kd_G0toS_*; the stringency of the G2/M checkpoint, 1/ *Kd_G2toG0_*. % Error (I), percentage of total mitotic error; Δ %Type (IV), absolute difference in the percentage of Type 1 and Type 2 cells. The white lines in (I) show contours. The white dashed lines in (IV) were manually drawn to emphasize the three regions with different arrest-phenotype composition: Type 2 dominant region (①), balanced region (②), and Type 1 dominant region (③). White X marks in (IV) indicate *Kd* combinations that yielded cell cycle profiles and arrest-failure rates similar to those observed in damaged RPE cells (see Fig. S6A). **(D)** Two testable perturbations of DNA damage checkpoints: inhibition of the G2/M checkpoint only or of both the G1/S and G2/M checkpoints, which can be achieved by ATR inhibition and p53 silencing, respectively. Dox, doxycycline; sh-p53, shRNA targeting p53; VE-821, a small molecule ATR inhibitor. **(E)** Percentage of micronucleated cells under damage with checkpoint perturbation (n = 4 biological replicates with >800 cells per condition). Cells were treated with VE-821(ATRi) 15 minutes before damage, treated with Doxycycline 7 hours before damage (pre damage sh-p53), or treated with Doxycycline 1 days after damage (post damage sh-p53). Cells were irradiated with 2Gy and subsequently cultured for three days before imaging. The post-damage culture time was shortened from 6 to 3 days to avoid overcrowding from checkpoint inhibition. **P < 0.01, ***P < 0.001, paired t test. **(F)** The effect of less stringent G2/M checkpoint on arrest-failure phenotype. Top: schematic of the checkpoint balance with or without ATRi. Bottom left: simulated effect of G2/M checkpoint inhibition on arrest-failure phenotypes in damaged cells. *Kd* backgrounds from cluster 1 in Fig. S6A were subjected to G2/M checkpoint perturbation (64-fold increase in *Kd_G2toG0_*) and subsequently exposed to their most vulnerable DNA damage levels. Result from the 3^rd^ model day after the induction of damage, each with three sets of sets of stochastic simulations with 500 seeding cells each. Values from the same *Kd* background are paired with a line. Bottom right: effect of ATRi on arrest-failure phenotypes in irradiated cells (n = 4 biological replicates with >800 cells per condition) – data from Fig. 6E. Cells were treated with or without VE-821 and irradiated with 2Gy and subsequently cultured for 3 days before imaging. Type 1 cells, 2C non-deformed micronucleated; Type 2 cells, ≥4C deformed micronucleated cells. Values from the same biological replicate are paired with a line. *P < 0.05, ***P < 0.001, paired t test. **(G)** The effect of more stringent G2/M checkpoint on arrest-failure phenotype. Top: schematic of the checkpoint balance in pre and post damage sh-p53. Bottom left: simulated effect of postponing G1/S & G2/M checkpoint inhibition on arrest-failure phenotypes in damaged cells. *Kd* backgrounds from cluster 1 in Fig. S6A were subjected to G1/S&G2/M checkpoint perturbation (64-fold increase in all the *Kd*s) before or one model day after the exposure to their most vulnerable DNA damage levels. Result from the 3^rd^ model day after the induction of damage, each with three sets of sets of stochastic simulations with 500 seeding cells each. Values from the same *Kd* background are paired with a line. Bottom right: effect of post damage p53 silencing on arrest-failure phenotypes in irradiated cells (n = 4 biological replicates with >800 cells per condition) – data from Fig. 6E. p53 was silenced 7 hours before or 1 days after 2Gy of irradiation. In both cases, cells were cultured for 3 days after irradiation before imaging. Type 1 cells, 2C non-deformed micronucleated; Type 2 cells, ≥4C deformed micronucleated cells. Values from the same biological replicate are paired with a line. **P < 0.01, ***P < 0.001, paired t test.

To examine the contribution of the checkpoints to arrest failure, we simulated arrest failure under different checkpoint stringencies (*Kds*) at a range of DNA damage levels. The result showed that, regardless of the stringencies, arrest failure was maximized under intermediate levels of DNA damage (Fig. 6B), consistent with our observations (Fig. 1C), suggesting overall vulnerability of the system to arrest failure under mild DNA damage. However, the maximum arrest failure varied greatly across different checkpoint stringencies (Fig. 6B), indicating that the extent of vulnerability can be tuned by *Kd*s. As expected, the system was most vulnerable when the checkpoint stringencies were the lowest, surprisingly however, a stronger stringency in a checkpoint did not necessarily lead to a lower arrest failure; rather, sensitive G1/S checkpoint and sub-maximal G2/M checkpoint did (Fig. 6C (I)). Since the arrest failure in Fig. 6C (I) is composed of both Type 1 and Type 2 errors, we simulated each type of error separately. The results showed that stringent G2/M checkpoint decreased the diploid Type 1 error but increased Type 2 error, whereas strong G1/S checkpoint reduced the fraction of Type 2 but increased that of Type 1 (Fig. 6C, (II) and (III), see more in *Discussion*), explaining the observed total arrest failure rates (Fig. 6C, (I)). To determine the checkpoint strength parameter regions in which each error type predominated, we plotted the differences in Type 1 and 2 error rates across the ranges of *Kd*s. This analysis revealed that the checkpoint landscape was divided into regions where one type dominated over the other (Fig. 6C, (IV), region ① and ③) or both types were in a relative balance (Fig. 6C, (IV), region ②), the latter of which contained the *Kd* regions that minimized total arrest failure. Together, the model predicted that strengthening one checkpoint can reduce one error type but adversely increases the other, resulting in an arrest-failure phenotypes tradeoff.

Cell cycle distribution and arrest-failure phenotypes in damaged RPE cells suggested that the cells fall in the “balanced” region (Fig. 6C, (IV), region ②, Fig. S6A), suggesting that the checkpoint system in non-cancerous cells may be optimized for the predicted tradeoff. To test the predicted tradeoff, we experimentally altered the checkpoint stringency and quantified the fraction of Type 1 and 2 RPE cells. We tested inhibition of the G2/M checkpoint only or inhibition of both checkpoints. Note that we could not specifically inhibit the G1/S checkpoint because p53 and ATM – established regulators of the G1/S checkpoint – also regulate the G2/M checkpoint^10,18,19^. Pretreating irradiated cells with VE-821 (Fig. 6D), a selective inhibitor of the G2/M checkpoint kinase ATR (Fig. S6B) ^16,37^, reduced 4C G0 cells – a byproduct of the activated G2/M checkpoint – and increased total micronucleated cells (Fig. 6E, Fig. S6D). Despite an increase in total micronucleation, VE-821 pretreatment only increased 2C-undeformed micronucleated cells but decreased 4C-deformed micronucleated cells (Fig. 6F), suggesting that a weaker G2/M checkpoint drives Type 1 error while attenuating Type 2 error, by allowing cells to complete mitosis rather than slipping into G0 without division. The model also predicted that inhibition of the G2/M checkpoint after DNA damage induction will become ineffective (Fig. S6E) because damaged G2 cells have already reset to G0. Therefore, delaying the inhibition of both checkpoints from before to after damage induction will decrease Type 1 error but increase Type 2 error if the tradeoff holds true (Fig. 6G). We found that p53 silencing 1 day post damage was indeed less effective in reducing the 4C G0 state compared to p53 silencing pre damage, despite efficient cell cycle promotion under damage (Fig. S6F). Both pre and post DNA damage p53 silencing increased total micronucleated cells (Fig. 6E). However, the arrest-failure phenotype in these two conditions was reversed as predicted; post (vs pre) damage p53 silencing resulted in decreased Type 1 error but exacerbated Type 2 error (Fig. 6G), by permitting cells time to bypass mitosis instead of forcing them through division with damage, further supporting the G2/M checkpoint’s anti-Type 1 but pro-Type 2 role. Together, these results demonstrate a critical tradeoff between diploid- and polyploid-derived arrest failures and explain why normal cells opt to evolve checkpoint strengths that balance these two error types.

## DISCUSSION

In this study, we identified two modes of arrest failure and the tradeoff between them, which is critical for minimizing total arrest failure rates. The Type 1-Type 2 tradeoff arises from the opposing effects of the G2/M and G1/S checkpoints on damage-induced polyploidy. The G2/M checkpoint initially halts cells at G2 via an ATM/ATR-dependent mechanism^10^ to allow repair of the damage. However, when timely repair fails, the G2/M checkpoint subsequently triggers mitotic skipping^8,9,38^ (Fig. 6A, “Reset”) via p53-driven p21 induction^13,39^, authorizing the first step toward endopolyploidy^8,9,38^. Thus, while the G2/M checkpoint prevents G2 cells from undergoing division with damage (blocking Type 1 cells), it also increases the subsequent chances of these cells becoming polyploid Type 2 cells. In contrast, the G1/S checkpoint prevents polyploid cells^7,9^ and thus the generation of Type 2 daughter cells by blocking the escape from G0 (Fig. 6A, “Escape”). However, because it prevents polyploid divisions, it indirectly increases the relative fraction of diploid cells – including Type 1 cells – in the population.

While a stronger tendency to arrest generally reduces the likelihood of arrest failure, it also hinders cell growth. As a result, cells require checkpoint mechanisms that are neither too permissive nor overly sensitive^40–43^. The Type 1-Type 2 tradeoff identified in this study uncovers a new dimension to this optimization problem; it shows that different risks need to be balanced with one another. Thus, the DNA damage checkpoint system must optimize checkpoint strengths considering multiple paths for failures. These tradeoffs, influenced by stochastic signaling pathways^25^, may explain why cells with functional checkpoints fail to arrest or escape arrest despite having unrepaired DNA damage^44^. Relatedly, in human cells with intact checkpoints, the G2/M checkpoint has a higher activation threshold than the G1/S checkpoint^22,26,28^. Our modeling suggests that this ‘negligent G2/M checkpoint’ may serve a functional purpose, since a less sensitive G2/M checkpoint, compared to G1/S checkpoint, is predicted to minimize overall arrest failure rates (Fig. 6C, (I)).

Earlier studies showed that G2 arrested cells gave rise to stress-induced endopolyploidy^7–9,14,15^. However, it was unclear whether these cells entered additional S phase via G1 or G0 since the commonly used FUCCI and related cell cycle reporters do not distinguish between G0 and G1^45^. Thus, in this study we probed the levels of endogenous Cdt1 and Geminin, both of which are known to be downregulated in G0^46^ (but not in G1) due to their transcriptional reliance on E2F activators^47^, as well as phosphorylated Rb, which marks cycling cells. Our data suggest that G0 is the predominant pathway, aligning with recent observations of cell state transitions from G2 to G0 under replication stress conditions^29^ or hypomitogenic stress^48^. Studies indicate that G0 induced by exogenous or endogenous^49^ DNA damage exhibits elevated G1 cyclin levels^50–52^ even though Rb is hypophosphorylated^29,53^. Consequently, such hyperaccumulation of G1 cyclins requires the G1/S checkpoint for the arrest of these damage-induced G0 cells^29^.

Our study suggests that polyploid division in stressed human cells with intact checkpoints may be more prevalent than commonly believed. Although cells with 8C DNA content were rare at a given time point due to their division, the footprint of polyploid mitosis became apparent through the sizable accumulation of lobulated 4C nuclei following mild DNA damage. These lobulated nuclei exhibited high chromosome missegregation rates (Fig. 4G), consistent with the observed erroneous nature of polyploid division across diverse contexts^33–35,54–57^. Recent studies using live-cell imaging similarly reported endoreplication and erroneous polyploid division^31^ in cells with intact checkpoints under various stress conditions^29–31^. These findings illustrate that polyploid division can result in chromosome missegregation in stressed normal cells, even without checkpoint defects caused by oncogene activation^9^ or compromised tumor suppressor functions^7,8^.

One question that emerges from our work is what happens to Type 1 and Type 2 cells in the long term. We speculate that most Type 1 and Type 2 cells will not be able to divide due to elevated p53^58,59^, aneuploidy^60,61^, or supernumerary centrosomes^62^. However, rare cells that manage to proliferate may give rise to oncogenic precursors, with Type 1 and Type 2 cells potentially leading to near-diploid and near-tetraploid cancers, respectively. Thus, the widespread occurrence of both near-diploid and near-tetraploid karyotypes in cancers may reflect the selection-driven tuning of the DNA damage checkpoint system to minimize total arrest failure rates amid the inherent Type 1-Type 2 tradeoff, thereby enhancing organismal fitness. Future studies are needed to clarify the cell fates of Type 1 and Type 2 cells and to explore whether we can direct cells toward specific routes to achieve a desirable therapeutic outcome.

## LIMITATION

Our study utilized a widely used non-cancerous human cell line, RPE1, and did not include other non-transformed and transformed human cell lines. The experimental perturbation of the G1/S checkpoint was not conducted since major checkpoint proteins with known function in the G1/S checkpoint also have roles in the G2/M checkpoint (e.g., p53^18,19^, ATM^17,18,63^, CHK2^64^). Additionally, the interplay between the DNA damage checkpoints and other potentially relevant checkpoints (e.g., the spindle assembly checkpoint) in arrest failure was not studied.

## RESOURCE AVAILABILITY

Requests for further information and codes should be directed to and will be fulfilled by the lead contact, Galit Lahav. This study did not generate new unique reagents. All original code has been deposited at GitHub [https://github.com/KotaroFuji/Markov_CellCycle_Checkpoints.git] and will be publicly available as of the date of publication. Any additional information required to reanalyze the data reported in this paper is available from the lead contact upon request.

## ACKNOWLEDGEMENTS

We thank Jose Ryes for valuable suggestions and sharing cell lines; members of the Lahav laboratory for comments, ideas, and support; Patrick J Flynn, Timothy J Mitchison, and Guang Yao for constructive discussion; the Nikon Imaging Center at HMS for support with immunofluorescence imaging. This research was supported by NIH grants R35 GM139572 and Ludwig Center at Harvard.

## AUTHOR CONTRIBUTIONS

Conceptualization, Methodology, Investigation, Software, Formal Analysis, Visualization, Writing – Original Draft, Review & Editing, K.F; Supervision, Project Administration, Writing – Review & Editing, A.J. and G.L; Funding Acquisition, G.L.

## STAR METHODS

### Key Resources Table

#### Reagent, Source, Identifier

##### Antibodies

Mouse monoclonal anti-H2A.X-P(Ser139), Millipore, JBW301; RRID: AB_310795

Rabbit monoclonal anti-CDT1, Cell Signaling Technology, D10F11; Cat#8064; RRID:AB_10896851

Mouse monoclonal anti-GMNN, DSHB, Cat#CPTC-GMNN-1; RRID:AB_2617263

Mouse monoclonal anti-p53, Santa Cruz Biotechnology, sc-126 (DO1); RRID: AB_628082

Rabbit monoclonal anti-phospho-Chk1(Ser345), Cell Signaling Technology, 133D3; Cat#2348 (also 2348S, 2348T, 2348L, 2348P); RRID:AB_331212

Rabbit monoclonal anti-phospho-Rb(Ser807/811), Cell Signaling Technology, Cat# 8516; RRID:AB_11178658

Rabbit polyclonal anti-gamma-Tubulin, Sigma-Aldrich, T5192; RRID: AB_261690

Mouse monoclonal anti-beta-Tubulin (E7), DSHB, Cat#E7

##### Chemicals, Peptides

Hoechst 33342, Invitrogen, Cat#H3570

5-ethynyl-2’-deoxyuridine (EdU), Invitrogen, Cat# C10338

Formamide (Deionized), Invitrogen, Cat# AM934

Blocking Reagent, Roche, 11096176001

Pan-centromere probe CENP-Cy5, PNA Bio, Cat#F3005

Doxorubicin (hydrochloride), Cayman Chemical, Item# 15007

Etoposide, Cayman Chemical, Item# 12092

Palbociclib, Sigma-Aldrich, PZ0383

Nutlin-3a, Sigma-Aldrich, SML0580

VE-821, Cayman Chemical, Item# 17587

BAY1217389, MCE, Cat# HY-12859

Barasertib (Synonyms: AZD1152), MCE, Cat# HY-10127

##### Bacterial and Virus strains

Tet -on-pLKO -p53shRNA (lentiviral), Reyes et al., 2018, N/A

pRRL-EF1a-mCerulean3-NLS (lentiviral), Reyes et al., 2018, N/A

CSII-EF-mCherry-hGeminin(1-110) (lentiviral), Jia Yun and S. Spencer, N/A

EF1a-iRFP-NLS, Jose Reyes, N/A

##### Experimental Models: Cell lines

Human: RPE-hTERT, S.J. Elledge Lab (Harvard Medical School), N/A

Human: RPE + tet-p53 shRNA + p53(shRNA-resistant)-DHFR(DD)-YFP, Reyes et al., 2018, N/A

Human: RPE + p53-mVenus + mCerulean3-NLS, Jose Reyes, N/A

Human: RPE + UbCp-p53-Venus + mCherry-Geminin + iRFP-NLS + tet-p53 shRNA, Jose Reyes, N/A

##### Software and Algorithms

P53Cinema Single Cell Tracking Software, Reyes et al., 2018, https://github.com/balvahal/p53CinemaManual

CellProfiler, Stirling et al., 2021, https://cellprofiler.org

ilastik, Berg et al., 2019, https://www.ilastik.org

Custom Python Script – Model, This work, Method S1, https://github.com/KotaroFuji/Markov_CellCycle_Checkpoints.git

## METHODS

### Cell Culture

Cells were passaged every 3 days and maintained at subconfluency in DMEM/F-12 medium supplemented with 10% fetal bovine serum (FBS). The medium was refreshed every 2 days in experiments spanning more than two days following the induction of DNA damage. For live-cell imaging, cells were switched to DMEM/F-12 + 5% FBS lacking phenol red and riboflavin to reduce autofluorescence. Cells were plated in fibronectin coated glass bottom 35mm dish or 12-well plate (MatTek Corporation) in low confluency (∼3K cells / 1cm^2^) a day before the induction of DNA damage.

### Induction of DNA Damage

To induce DNA damage, cells were exposed to X-ray (RS-1800 irradiator, Radsource), Doxorubicin, or Etoposide. In Doxorubicin or Etoposide treatment, the medium containing the drug was removed two days following the initial exposure, and cells were washed twice and being replaced with the fresh medium to allow recovery from the drug treatment.

### Immunofluorescence

Cells were washed once with PBS and subsequently fixed with 4% paraformaldehyde for 12min at room temperature or ice-cold methanol for 3min on ice. Cells were washed twice with PBS and subsequently blocked/permeabilized with 0.1% Triton X-100 and 2% BSA in PBS for 30min at room temperature. Cells were incubated overnight with primary antibody at 4 °C. Cells were washed thrice with PBS, and subsequently incubated with secondary antibody at room temperature for 1hr. Finally, cells were washed twice with PBS before Hoechst 33342 staining (1 μg/ml), after which the cells were further washed thrice with PBS. Cells were imaged using Nikon Eclipse TE-2000 inverted microscope or Yokogawa CSU-X1 spinning disk confocal microscope, with a 20X or 40X Plan Apo objective and a Hammamatsu Orca ER camera. Images were taken in Z stacks, each with 17 slices with a 1μm step size.

### EdU Assay

Cells were incubated with EdU (10μM in the medium) for 20min before washing, fixation, and blocking/permeabilization as described above. Cells were incubated with the Click-iT reaction cocktail (2mM CuSO4, 8μM sulfo-Cyanine3 azide, and 10mg/ml L-Ascorbic acid in PBS) for 30min at room temperature. Cells were washed once with PBS before Hoechst 33342 staining (1 μg/ml). When combined with immunofluorescence, the incubation with the Click-iT reaction cocktail was performed after the first washing step following secondary antibody staining. Cells were imaged using Nikon Eclipse TE-2000 inverted microscope, with a 20X or 40X Plan Apo objective and a Hammamatsu Orca ER camera. Images were taken in Z stacks, each with 17 slices with a 1μm step size.

### Live-cell Microscopy

Cells were imaged using a Nikon Eclipse TE-2000 inverted microscope with a 20X Plan Apo objective and a Hammamatsu Orca ER camera, with an environmental chamber controlling temperature, humidity, and carbon dioxide (5%). The images were acquired every 15min using the MetaMorph Software.

### Checkpoint Inhibitions and Cell-cycle Perturbations

To silence p53 before or after the induction of DNA damage, cells stably expressing doxycycline inducible shRNA targeting p53 were exposed to 500ng/ml of doxycycline ∼7hr before or one day after the induction of DNA damage, respectively. To inhibit ATR before the induction of DNA damage, cells were incubated with VE-821 15min before the induction of DNA damage. To pre-arrest cells, cells were cultured with Palbociclib or Nutlin-3a one day before the induction of DNA damage; the drug was kept in the medium after the induction of DNA damage if cell-cycle arrest was to be maintained. Medium containing one of these drugs was replaced every other day when experiments spanned longer than two days.

### PNA FISH Assay

Cells were washed once with PBS and subsequently fixed in an ice-cold fixative (3:1 ratio of methanol and acetic acid) for 5min at 4 °C. Cells were washed four times with PBS and subsequently underwent quick serial dehydration using 70%, 85%, and 100% ethanol, with cells immersed within for two, one, and one minute(s), respectively. Cells were airdried overnight at room temperature. Cells were incubated with pre-heated hybridization buffer containing PNA probe (0.2 μM PNA probe, 20mM Tris pH 7.4, 60% formamide, and 0.5% blocking reagent) for 10min at 80 °C and subsequently for 2hr at room temperature, with controlled humidity to prevent drying. Cells were washed thrice in washing solution (0.1% Tween-20 in 2X SSC); twice with each 10min incubation at 55°C, and lastly once at room temperature. Cells underwent Hoechst 33342 staining (1 μg/ml) and were subsequently washed with 2X SSC for 2min, 1X SSC for 2min, and finally with H_2_O for 2min. Cells were put in PBS and imaged using Nikon Eclipse TE-2000 inverted microscope with a 40X Plan Apo objective and a Hammamatsu Orca ER camera. Images were taken in Z stacks, each with 17 slices with a 1μm step size.

### Image Quantification and Analysis

#### Quantification of Nuclear Morphology

The degree of nuclear deformation was quantified by irregularity score, which is obtained by

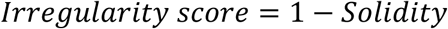

And solidity of a nucleus is defined by

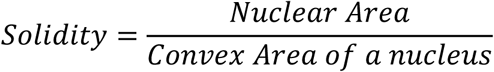

Solidity was calculated using the MeasureObjectSizeShape function in CellProfiler^67^, based on automatically segmented primary nuclei as described in the nuclear segmentation section below. Deformed nuclei under DNA-damage conditions were defined as those whose irregularity scores exceeded the top 2 percentile of irregularity scores in batch-controlled non-perturbed cells. Other nuclear features related to size and shape were also quantified using the same MeasureObjectSizeShape function. Micronuclei were identified and quantified as described in the nuclear segmentation section.

#### Pixel Classification and Nuclear Segmentation

To segment images, the max projection images of Hoechst 33342 staining were first processed by a machine-learning based pixel classifier in ilastik^68^. We used three of the center Z slices (out of 17) for max projection to enhance sharpness while avoiding out-of-focus light. The algorithm was pre-trained to classify each pixel into a background, primary nucleus, or micronucleus. The algorithm outputted each Hoechst 33342 staining image (Fig. S7A, (I)) as a colored probability map reflecting the probabilities of each pixel to be a background (red), primary nucleus (green), or micronucleus (blue) (Fig. S7A, (II)). These probability maps were fed into CellProfiler^67^ for the segmentation of primary nuclei using the Minimum Cross-Entropy thresholding method with the adaptive (as opposed to global) strategy provided in the software (Fig. S7A, (III)). Any object whose diameter fell outside the set pixel range was discarded (25 ∼ 200 pixels for an image taken with an 20X objective without pixel binning; the pixel range was multiplied according to the magnification and pixel binning settings). We did not de-clump objects during this segmentation, as this may lead to a deformed nuclei being falsely recognized as a clumped object. Seeding cells at low confluency was used to circumvent cell crowding. After nuclear segmentation, peripheral region for each primary nucleus was determined by expanding each nucleus by certain pixels (100 pixels for an image taken using 20X objective without binning; the pixel number was multiplied according to the magnification and pixel binning settings; Fig. S7A, (IV)) and subsequently masking the nuclear region from such expanded area (Fig. S7A, (V)). Micronuclei within nuclear peripheral regions were segmented based on probability map (Fig. S7A, (VI), (II)). Any micronuclei that were outside of or touched the border of these nuclear peripheral regions, or whose diameter fell outside a certain range (3 ∼ 35 pixels for an image taken using 20X objective without binning; the pixel number was multiplied according to the magnification and pixel binning settings), were excluded. Finally, each micronucleus was paired with its parental primary nucleus, and micronuclei number was quantified for each primary nucleus using the RelateObjects function in CellProfiler^67^.

#### Feature Extraction/Quantification from Images

Features from immunostaining or FISH images were extracted using CellProfiler^67^ and ilastik^68^. Max and sum projection images were corrected for global/systematic illumination bias using the Fit Polynomial method in the CorrectIllumination functions in CellProfiler^67^. Since all antibodies we quantified targeted nuclear proteins, we subtracted minimum intensity in each nucleus’ peripheral region (defined as in Fig. S7A, (V)) as a background signal from nuclear signal. Indices of signal intensity and heterogeneity within the segmented areas were quantified using the MeasureObjectIntensity and MeasureGranularity functions in CellProfiler^67^. To detect γ-H2AX or γ-Tubulin foci, we referenced the Speckle Counting analysis shared in <u>CellProfiler’s example</u> <u>pipelines</u>; we enhanced foci features using the EnhanceOrSuppressFeatures function and counted these foci within each primary nucleus using the Robust Background method with the global strategy. To quantify centromere, we trained Ilastik^68^, similar to what was described in the segmentation section above, and processed probability map in Cellprofiler^67^ to identify centromeres specifically within the segmented primary nuclei and micronuclei.

#### Cleaning Data

The purpose of cleaning data was to filter out unwanted objects identified in the segmentation process described above. For example, micronuclei and mitotic chromosomes can be segmented as primary nuclei. We opted gating strategies similar to Flow Cytometry analyses to filter out micronuclei and mitotic chromosomes falsely segmented as primary nuclei, using the R package gatepoints^69^. We gated objects based on DNA content and nuclear area, which allowed us to easily identify and gate out objects with sub-G1 DNA content such as micronuclei (Fig. S7B). We also gated out mitotic chromosomes, leveraging on their compaction with ultra-high DNA staining. We calculated mean DNA intensity divided by area of DNA staining (i.e., “nuclear area” when segmented), and filtered out objects with exceptionally high such values (Fig. S7C).

#### Trajectory Inference

To infer polyploid trajectory following irradiation, we used the R package slingshot^36^ and implemented their analysis flow presented in their vignette. Images stained for DNA, Cdt1, Geminin, and EdU were segmented, quantified, and cleaned as described above. The quantified features were centered and scaled using the scale function in R^70^, assuming that the extracted features were all relevant in deducing polyploidization. To deduce the trajectory from G2 cells to undivided polyploid cells, we specifically analyzed non-micronucleated cells with ≥4C DNA content for trajectory inference. We performed dimensionality reduction using uniform manifold approximation and projection and clustered cells using *k*-means method, with a *k* of 4. We then constructed a minimum spanning tree on the identified clusters by applying the getLineages function in the package. The starting cluster was set to the one enriched with cells before irradiation, and the end cluster was set to the one enriched in polyploid cells with ≥8C DNA content. Finally, pseudotime was obtained using the getCurves and slingPseudotime functions. The obtained pseudotime and other quantified features were visualized as tSNE plot using the R package Rtsne^71^.

#### Single-cell Tracking

We used the previously published P53Cinema Single Cell Tracking Software^25^ for the tracking live-cell imaging data. We randomly tracked cells that divided following irradiation using NLS reporter signals.

#### Data Formatting and Visualization

Data was formatted using custom python and R scripts and visualized using the R package ggplot2^72^.

#### Magnification

To determine the minimal magnification required for accurate quantification of irregularity scores in RPE cells, we imaged multiple nuclei with varying levels of nuclear deformation using 100X objective. We serially decreased the pixel resolution of these images *in silico* (Fig. S7D), to mimic images taken by lower magnifications. Irregularity score was calculated for each nucleus under different pixel resolution (Fig. S7E). As the pixel resolution decreased, the dynamic range of irregularity scores shrank especially under around 36 pixels (Fig. S7E), suggesting that the quantification becomes inaccurate when imaged nuclei display diameters below this threshold. Since most nuclei imaged with 20X objective without pixel binning displayed diameter larger than 36 pixels (Fig. S7F), we used at least 20X magnification for our imaging procedures.

#### Micronuclear centromeres as markers of chromosome missegregation

We experimentally validated that micronuclei generated from acentric chromatin fragments or chromosome missegregation can be differentiated by the presence or absence of centromeres in micronuclei; whole (or near whole) chromosome missegregation frequently generates micronuclei containing centromeres, whereas micronuclei generated from damaged DNA lack centromeres (Fig. S4C). Centromeres within micronuclei were quantified using PNA FISH for CENP-B box DNA repeats. Cells treated with irradiation and the ATR inhibitor, which elevated micronucleation by promoting mitosis with DNA damage (Fig. S8A-D), generated micronuclei lacking centromeres (Fig. S8E). Conversely, both the MPS1 inhibitor BAY-1217389 and the AURKB inhibitor Barasertib (antimitotics that disrupt chromosome segregation), generated micronuclei positive for centromeres without elevating γ-H2AX foci (Fig. S8A-E). Thus, we concluded that centromere detection in micronuclei using PNA FISH informs the origin of micronuclei.

#### Simulation

The aim of our simulation was to develop an intuitive understanding of what leads to mitotic errors in DNA damaged cells. We constructed a stochastic Markov chain-like model, in which cells transition from one cell cycle state to another based on certain probabilities, which DNA damage and DNA damage checkpoint could affect. The table below shows the probabilities of cells moving from one state to another.

| From \ To | G0 | G1 | S | G2 | M |
| --- | --- | --- | --- | --- | --- |
| G0 | $1-P_{G0\vert S}$ | 0 | $P_{G0\vert S}$ | 0 | 0 |
| G1 | $P_{G1\vert G0}$ | $1-P_{G1\vert G0}-P_{G1\vert S}$ | $P_{G1\vert S}$ | 0 | 0 |
| S | 0 | 0 | $1-P_{S\vert G2}$ | $P_{S\vert G2}$ | 0 |
| G2 | $P_{G2\vert G0}$ | 0 | 0 | $1-P_{G2\vert G0}-P_{G2\vert M}$ | $P_{G2\vert M}$ |
| M | 0 | $P_{M\vert G1}$ | 0 | 0 | $1-P_{M\vert G1}$ |

Probabilities are defined as

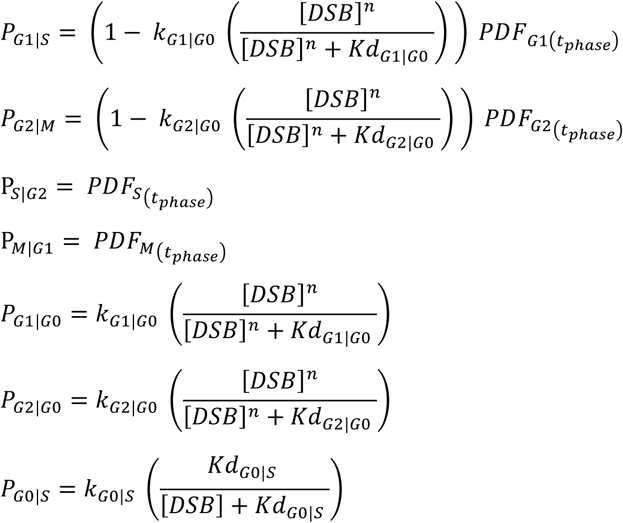

Un-perturbed cells in the cell cycle (i.e., cells in G1, S, G2, or M phase) stochastically transition to the next phase of the cell cycle based on the probability density function at the corresponding phase, denoted by *PDF*, and the time they spent in that particular phase, *t_phase_*. The probability density functions were computationally generated based on live-cell imaging data that quantified the duration of each cell-cycle phase in unperturbed RPE cells, as specified in Table S1. We assumed that DNA damage, denoted by [*DSB*], above a certain threshold, determined by the *Kd* values, can drive G1 or G2 cells into G0 with rapid switch-like property. We set a hill coefficient *n* of 3, which confers a physiologically relevant level of switch-like property to the system. DNA damage also prevents G0 cells from escaping G0 and reentering the cell cycle. Based on the leakiness of the G1/S cell-cycle maintenance^6,25^, we assumed *n* of 1 for the arrest maintenance.

We assumed that the incurred [*DSB*] gets repaired in constant rate over time.

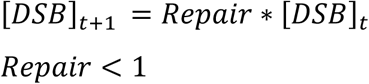

We simulated mitotic error according to the level of DNA damage and their ploidy state upon mitosis. The probability of harboring mitotic error in a cell that just underwent division was set as

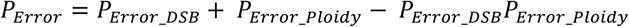

In which

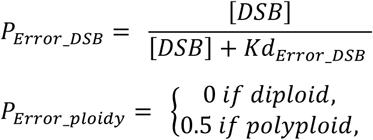

Here, the probability of mitotic error depends on both DNA damage and ploidy state. Specifically, having DNA damage or being polyploid increases the likelihood of mitotic error, specified accordingly by *P_Error_DSB_* or *P_Error_ploidy_*. *Kd_Error_DSB_* was arbitrarily set to 1 as any number will do. *P_Error_ploidy_* in polyploid daughters was set to 0.5 based on our observation that majority of 8C mothers give rise to deformed nuclei. We assumed that polyploid division contributes to mitotic error independent of DNA damage level, consistent with erroneous nature of polyploid mitosis^33–35,54–57^. We defined diploid and polyploid daughters with mitotic error as Type 1 and Type 2 cells, respectively.

---

**Supplementary Table 1.**

| Parameter | Definition | Value | Units | Source |
| --- | --- | --- | --- | --- |
| $*PDF_{G1}$ | Probability density function for G1, determined by the mean ($M$) and standard deviation ($SD$) of the G1 phase duration | $(M, SD) = (560\text{min}, 130\text{min})$ | | From this work |
| $*PDF_S$ | Probability density function for S, determined by the mean ($M$) and standard deviation ($SD$) of the S phase duration | $(M, SD) = (200\text{min}, 90\text{min})$ | | From this work |
| $*PDF_{G2}$ | Probability density function for G2, determined by the mean ($M$) and standard deviation ($SD$) of the G2 phase duration | $(M, SD) = (220\text{min}, 60\text{min})$ | | From this work |
| $*PDF_M$ | Probability density function for M, determined by the mean ($M$) and standard deviation ($SD$) of the M phase duration | $(M, SD) = (30\text{min}, 10\text{min})$ | | Araujo et al., 2016<sup>1</sup> |
| $n$ | hill coefficient for DNA damage checkpoint activation | 3 | | Inferred based on the switch-like effect of DNA damage checkpoint on arresting cells |
| $Repair\ rate$ | Long-term repair rate of DNA damage | 0.9982279 | $10\text{min}^{-1}$ | From this work |
| $k_{G0\vert S}$ | Maximum likelihood a cell in G0 reenters the cell cycle within 10min | 0.005192669 | $10\text{min}^{-1}$ | Inferred based on Lo et al., 2020<sup>2</sup> |
| $k_{G1\vert G0}$ | Maximum likelihood a cell in G1 initiates the arrest within 10min | 0.174595815 | $10\text{min}^{-1}$ | Inferred based on Deckbar et al., 2011<sup>3</sup> |
| $k_{G2\vert G0}$ | Maximum likelihood a cell in G2 initiates the arrest within 10min | 0.99 | $10\text{min}^{-1}$ | Inferred based on Deckbar et al., 2011<sup>3</sup> |

\*PDF was calculated based on 10,000 randomly-generated in silico cells with the statistical distribution with the specified M and SD

## SUPPLEMENTARY FIGURE LEGENDS

**Figure S1:**
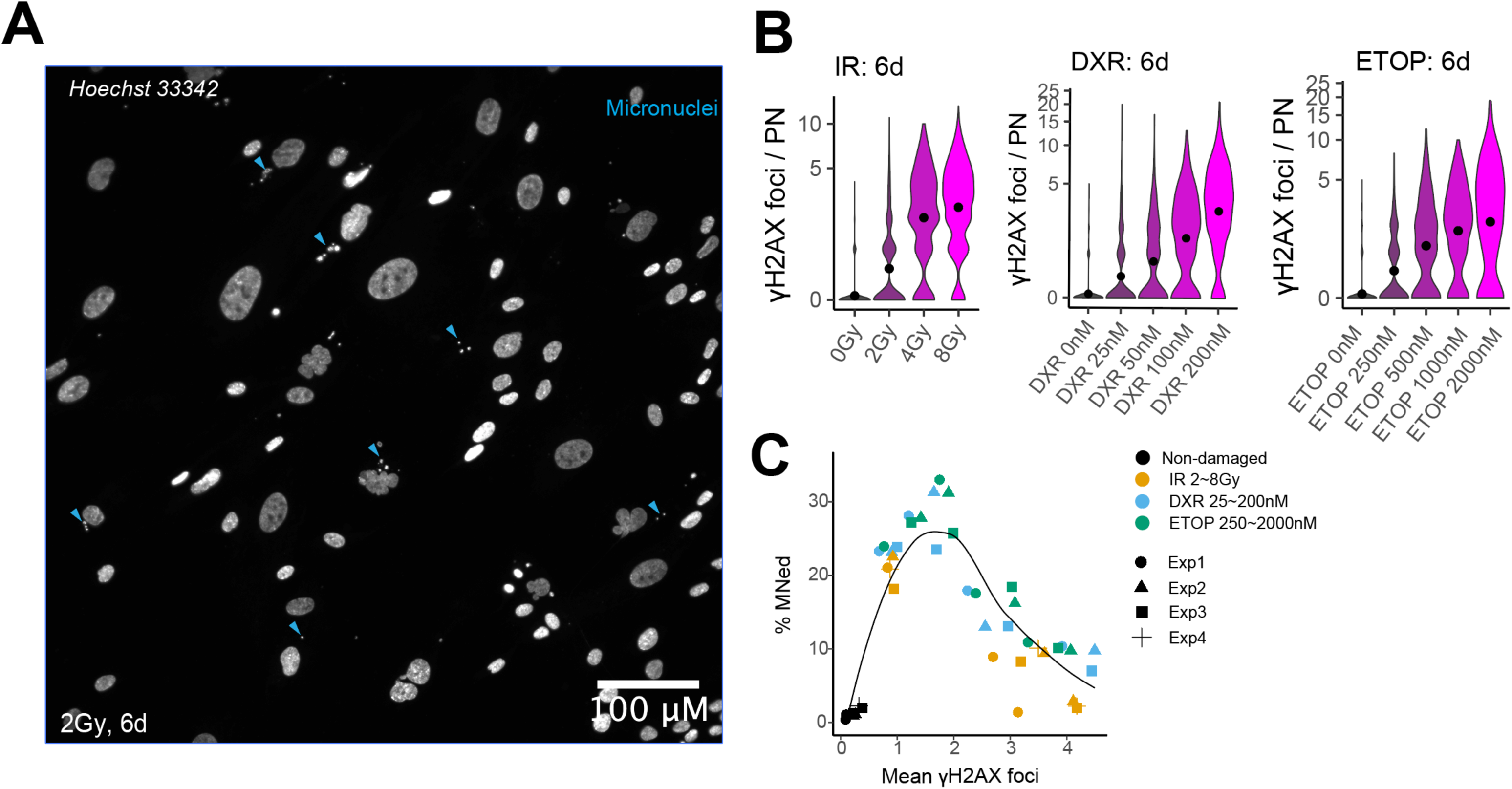
Intermediate levels of DNA damage maximize the fraction of micronucleation. **(A)** A field view of 20x image taken from cells irradiated with 2Gy and subsequently cultured for 6 days. Examples of micronuclei are indicated by the colored arrowheads. **(B)** Number of γ-H2AX foci per primary nucleus in cells treated with different doses of various DNA-damaging agents (n > 1000 cells per dose). Irradiated cells were cultured for 6 days. Cells were exposed to one of the topoisomerase inhibitors for 2 days then recovered for 4 days before imaging. IR, irradiation; DXR, Doxorubicin; ETOP, Etoposide. Foci with clear and strong staining were quantified. PN, primary nucleus. The dots represent means. **(C)** Quantification of the mean γ-H2AX foci and the percentage of micronucleated cells in cells treated with different doses of various DNA-damaging agents. Each point represents a population. Quantification from ≥3 biological replicates with >250 cells per population – Data from Fig. 1C. Shape symbols indicate independent biological replicates.

**Figure S2:**
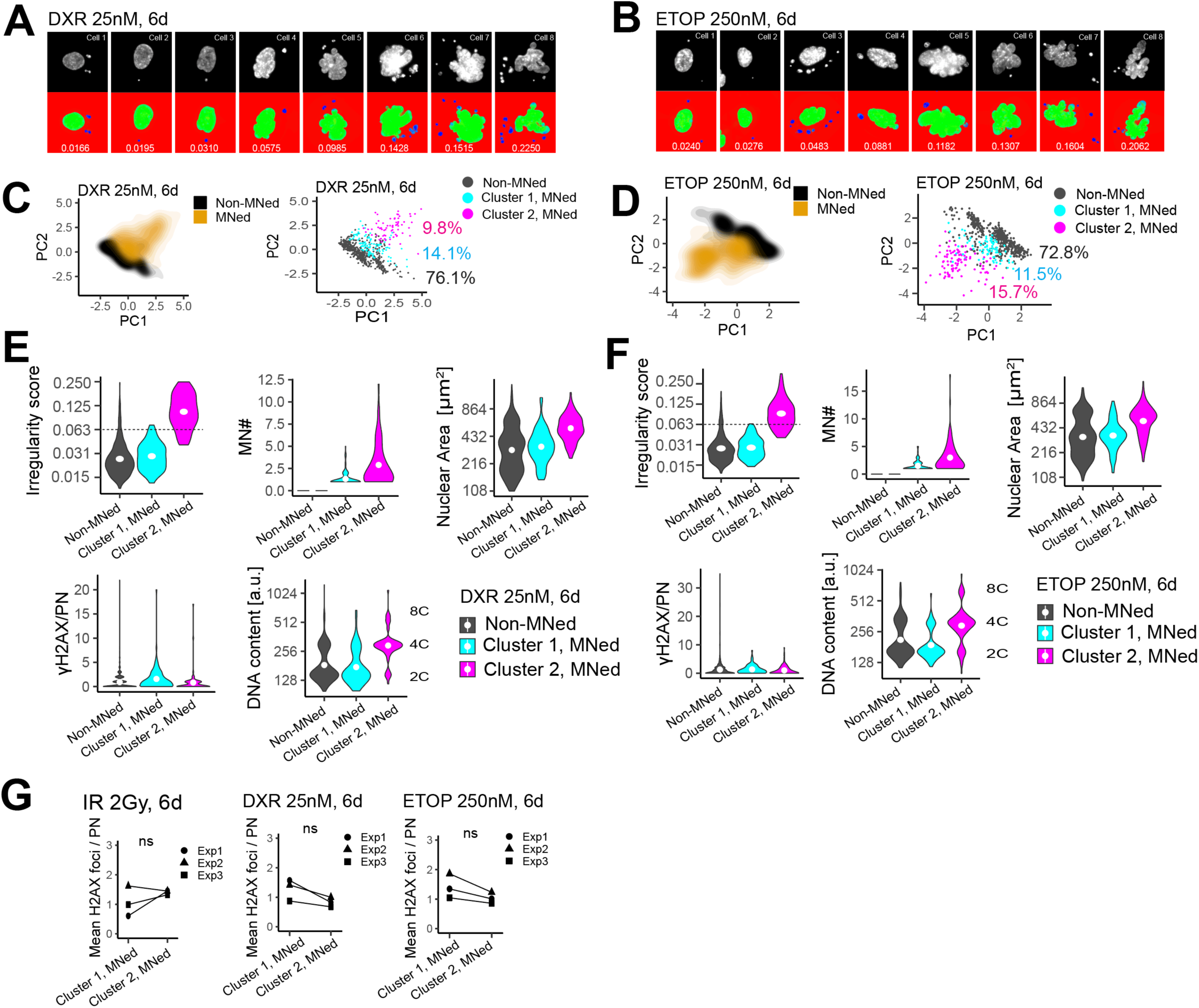
Two distinct arrest-failure phenotypes in DNA-damaged cells. **(A, B)** Nuclear images of micronucleated cells. DXR, Doxorubicin; ETOP, Etoposide. **(C, D)** 2D Scatterplot of PC1 and PC2 (left) and the two clusters within the micronucleated subset (bright) (n > 650 cells). Cells were treated as in A or B. **(E, F)** The distribution of the five nuclear features in micronucleated cluster 1 and cluster 2 cells (n > 650 cells) – data from C or D. **(G)** The mean γ-H2AX foci per primary nucleus in micronucleated cluster 1 and cluster 2 cells (n = 3 biological replicates with ≥140 micronucleated cells each). Values from the same biological replicates are paired with a line. Shape symbols indicate independent biological replicates. ns, P>0.05 in paired t test.

**Figure S3:**
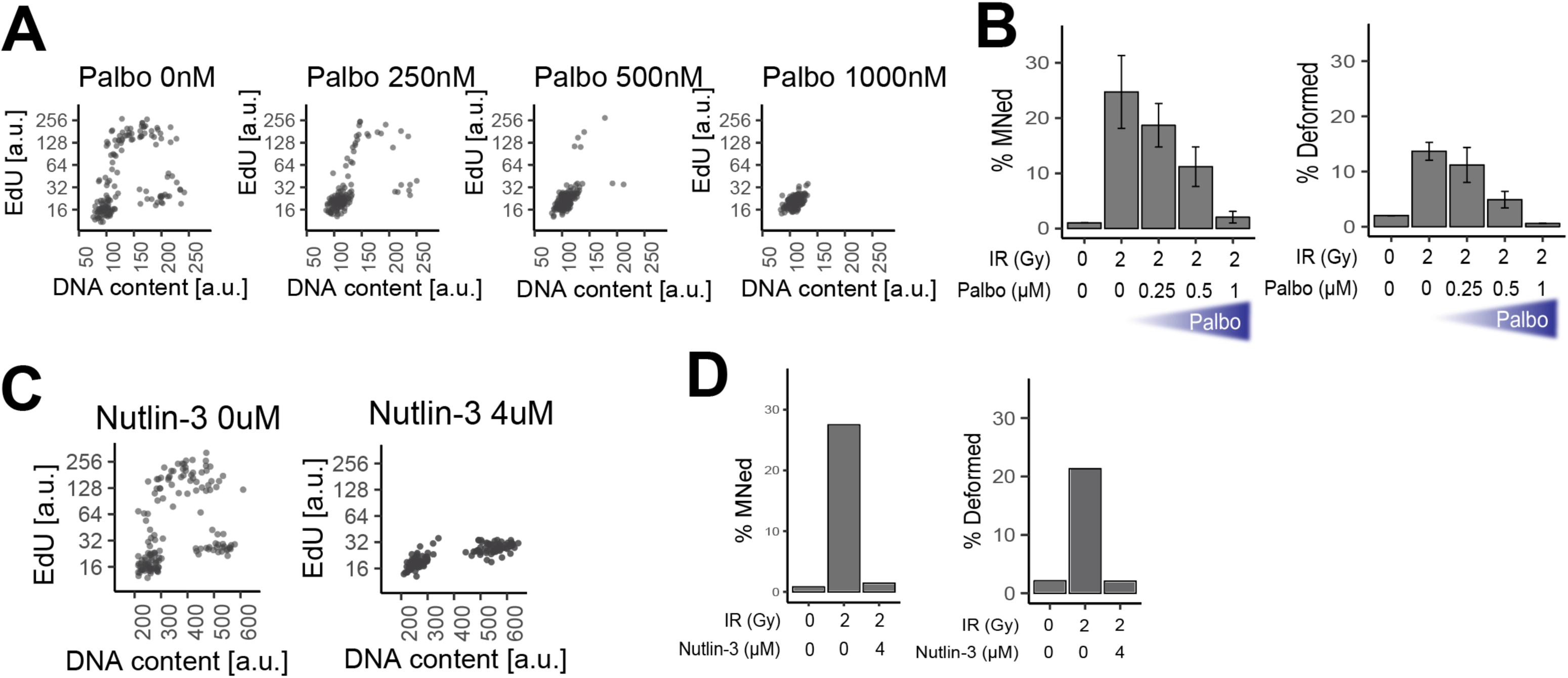
Nuclear deformation requires division. **(A)** Profile of DNA content and EdU signal in cells treated with different concentrations of Palbocyclib for one day, quantified by microscopy (n = 150 cells per condition). Palbo, Palbocyclib. **(B)** Percentage of micronucleation and nuclear deformation in non-damaged and irradiated cells pre-arrested and co-treated with increasing concentrations of Palbocyclib (n = 2 biological replicates with >300 cells each). Cells were pre-arrested by Palbocyclib for a day, irradiated, and subsequently cultured for 6 days in the continued presence of Palbocyclib before imaging. Palbo, Palbocyclib. Error bar, SEM. **(C)** Profile of DNA content and EdU signal in cells treated with or without Nutlin-3 for one day and quantified by microscopy (n = 150 cells per condition). **(D)** Percentage of micronucleation and nuclear deformation in non-damaged cells and irradiated cells with or without Nutlin-3 treatment (n > 800 cells per condition). Cells were pre-arrested by Nutlin-3 for a day, irradiated, and subsequently cultured for 6 days in the continued presence of Nutlin-3 before imaging.

**Figure S4:**
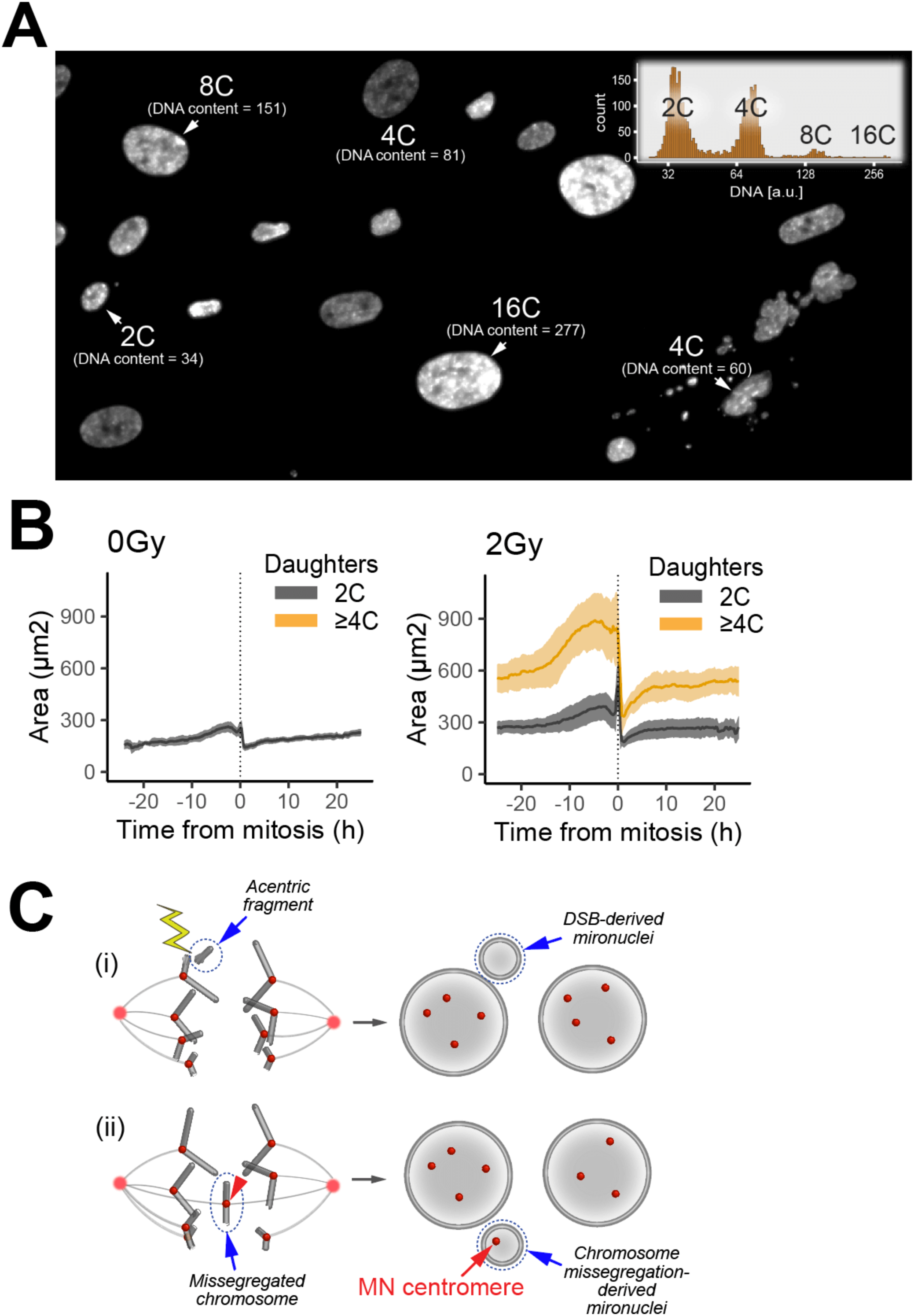
Damaged-induced polyploidy in normal human cells. **(A)** Examples of DNA-stained irradiated cells, including polyploid nuclei (8C and 16C). (Inset) Histogram of DNA content from cells in A (n > 3000 cells) **(B)** Quantification of nuclear areas in 4C and 8C mothers and their daughters (n = 82, 237, and 137 cells correspondingly for non-damaged 4C, irradiated 4C, and irradiated 8C subsets) – data from Fig. 4E. The cells were synchronized *in silico* according to their mitotic timing. The median and interquartile range are respectively shown in thick lines and colored ribbons. **(C)** Schematics of two distinct scenarios that generate micronuclei through DSBs or chromosome missegregation. DSBs can generate acentric DNA fragments, which prevent them from being properly segregated during division, resulting in micronuclei that lack centromeres (i). Chromosome missegregation can manifest after division as micronuclei with centromere(s) contained within (ii). MN, micronuclei.

**Figure S5:**
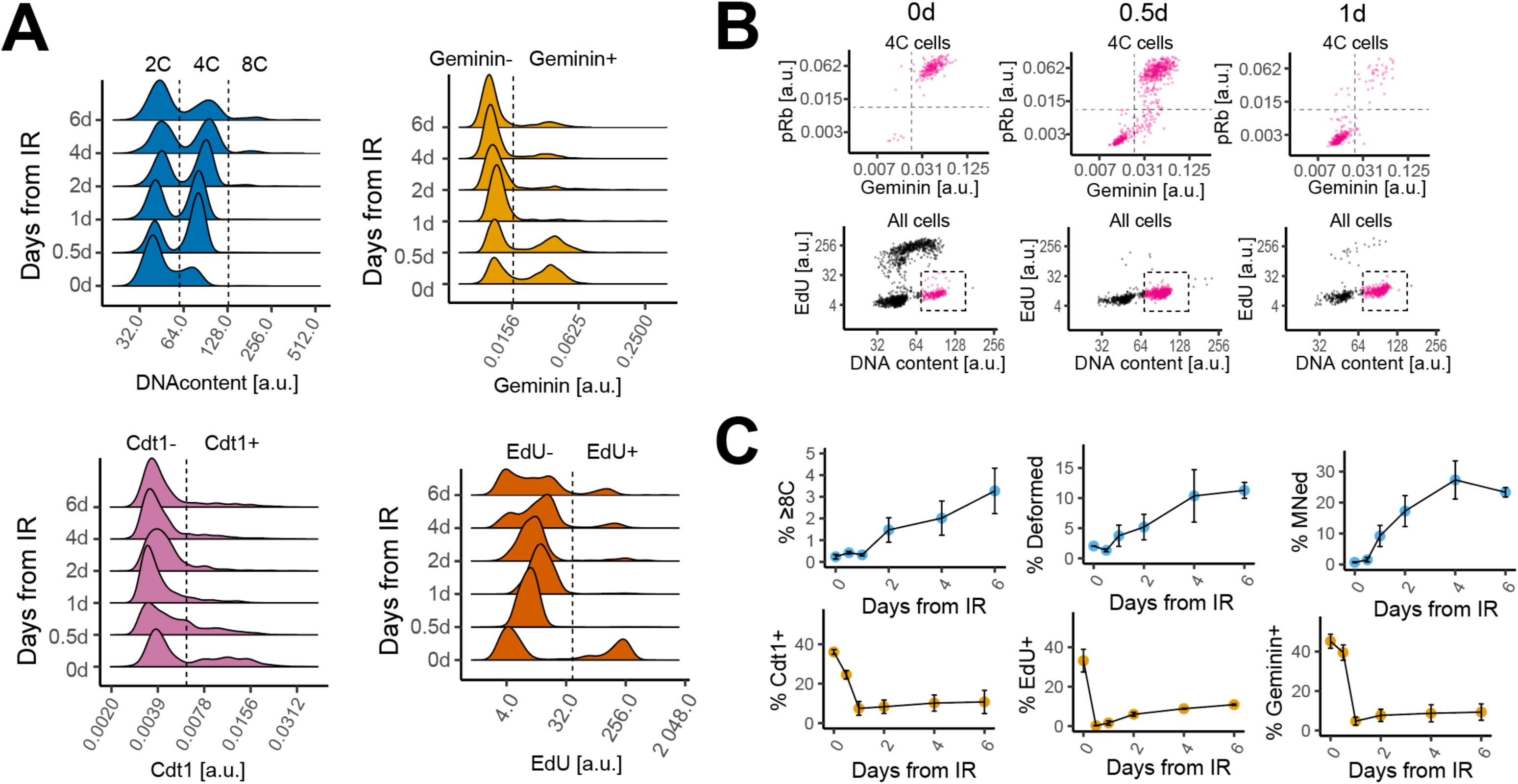
G2-to-G0 transition precedes irradiation-induced polyploidization. **(A)** Distribution of DNA content, Cdt1, Geminin, and EdU signal in cells exposed to 2Gy irradiation and cultured for up to 6 days (n > 570 cells per timepoint) – data from Fig. 5C. Dashed lines indicate thresholds for DNA content or marker positivity classification. **(B)** Quantification of phosphorylated Rb and Geminin specifically in cells with 4C DNA content (magenta), at the indicated timepoints following 2Gy irradiation (n > 690 cells per timepoint). Pulse-labeled EdU, Geminin, and phosphorylated Rb were quantified in single cells by Click-iT chemistry and immunofluorescence. Cells with 4C DNA content were gated as shown in the bottom row, using DNA staining and EdU signal. **(C)** Percentage of cells exhibiting the indicated feature at multiple timepoints following 2Gy irradiation (n = 2 biological replicates with >570 cells each). ≥8C: primary nuclei with DNA content equal or greater than 8C. Error bar, SEM.

**Figure S6:**
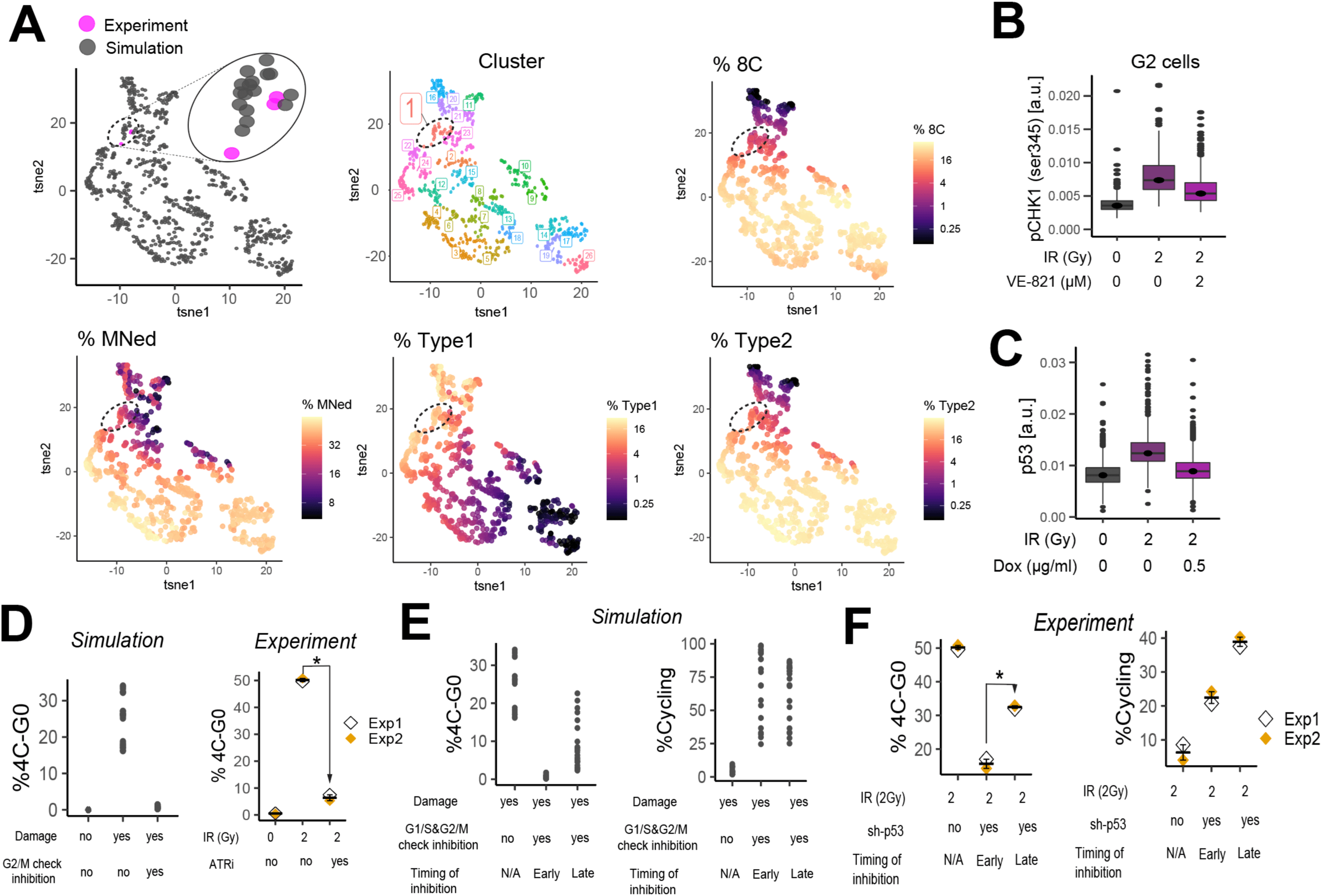
Effect of checkpoint stringencies on arrest-failure types and the cell cycle. **(A)** Profile of cell cycle state and arrest-failure phenotype under damage in simulations and experiments. Each dot in the t-SNE plots represents a population of cells from simulations or experiments. Simulation: each *Kd* value was varied between 0.001 and 64, and 729 unique *Kd* combination were simulated under their most vulnerable DNA damage level – simulations from Fig. 6C. Result from the 6^th^ model day after the induction of damage. Each condition with three sets of sets of stochastic simulations with 100 seeding cells each. Experiment: cells were exposed to 2Gy of irradiation and subsequently cultured for 6 days (n = 3 biological replicates with >600 cells per replicate) – data from Fig. 1C. For each population (whether from simulations or experiments), the fractions of 2C, 4C, 8C, Type 1, and Type 2 cells were quantified. These fractions were used for t-SNE analysis, the result of which was used for hierarchical clustering. The *Kd* combinations in cluster 1 (the ones in a dashed oval) were used for the simulation results in Fig. S6D-F and Fig. 6F, G. **(B)** The effect of VE-821 on serine 345 phosphorylation on pCHK1 (n > 1400 cells per condition). The cells were pretreated with VE-821 prior to 2Gy of irradiation and subsequently cultured for 2 hours in the continued presence of the drug before immunostaining. Cells with G2 DNA content were analyzed. The boxes and the lines represent interquartile ranges and medians, respectively. **(C)** The effect of the inducible shRNA targeting p53 on p53 levels (n > 1400 cells per condition). The cells were treated with doxycycline before their exposure to 2Gy of irradiation and subsequently cultured for 2 hours in the continued presence of the drug before p53 immunostaining. The boxes and the lines represent interquartile ranges and medians, respectively. **(D)** The effect of the G2/M checkpoint perturbation on the 4C G0 state. Left: simulated effect of the G2/M checkpoint inhibition on the 4C G0 state. Result from the 3^rd^ model day after the induction of damage in background parameter conditions found in cluster 1 of Fig. S6A. Percentage of 4C G0 cells was calculated with or without 64-fold increase in *Kd_G2toG0_*. Three sets of stochastic simulations were run for each condition (500 seeding cells each). Right: effect of VE-821 treatment on the 4C G0 state (n = 2 biological replicates with >1400 cells). Cells were treated with VE-821 and exposed to 2Gy of irradiation and subsequently cultured for 3 days before imaging. Cells with 4C G0 state were defined as ones with non-phosphorylated Rb and 4C DNA content. Error bar, SEM. *P < 0.05, one-tailed paired t test. **(E)** Simulated effect of postponing the G1/S&G2/M checkpoint inhibition under damage on the cell cycle. Result from the 3^rd^ model day in background parameter conditions found in cluster 1 of Fig. S6A with perturbed *Kd*s. *Kd*s were increased 64-fold from their background states just before or one model day after the damage. Percentage of 4C G0 cells and cycling cells was calculated from three sets of stochastic simulations (500 seeding cells each). **(F)** Effect of postponing p53 silencing under damage on the cell cycle (n = 2 biological replicates with >1400 cells). p53 was silenced 7 hours before or one day after 2Gy of irradiation. Cells were cultured for 3 days after irradiation before being imaged. Cells with 4C G0 state were defined as ones with non-phosphorylated Rb and 4C DNA content. Cycling cells were defined as ones with phosphorylated Rb. Error bar, SEM. *P < 0.05, one-tailed paired t test.

**Figure S7:**
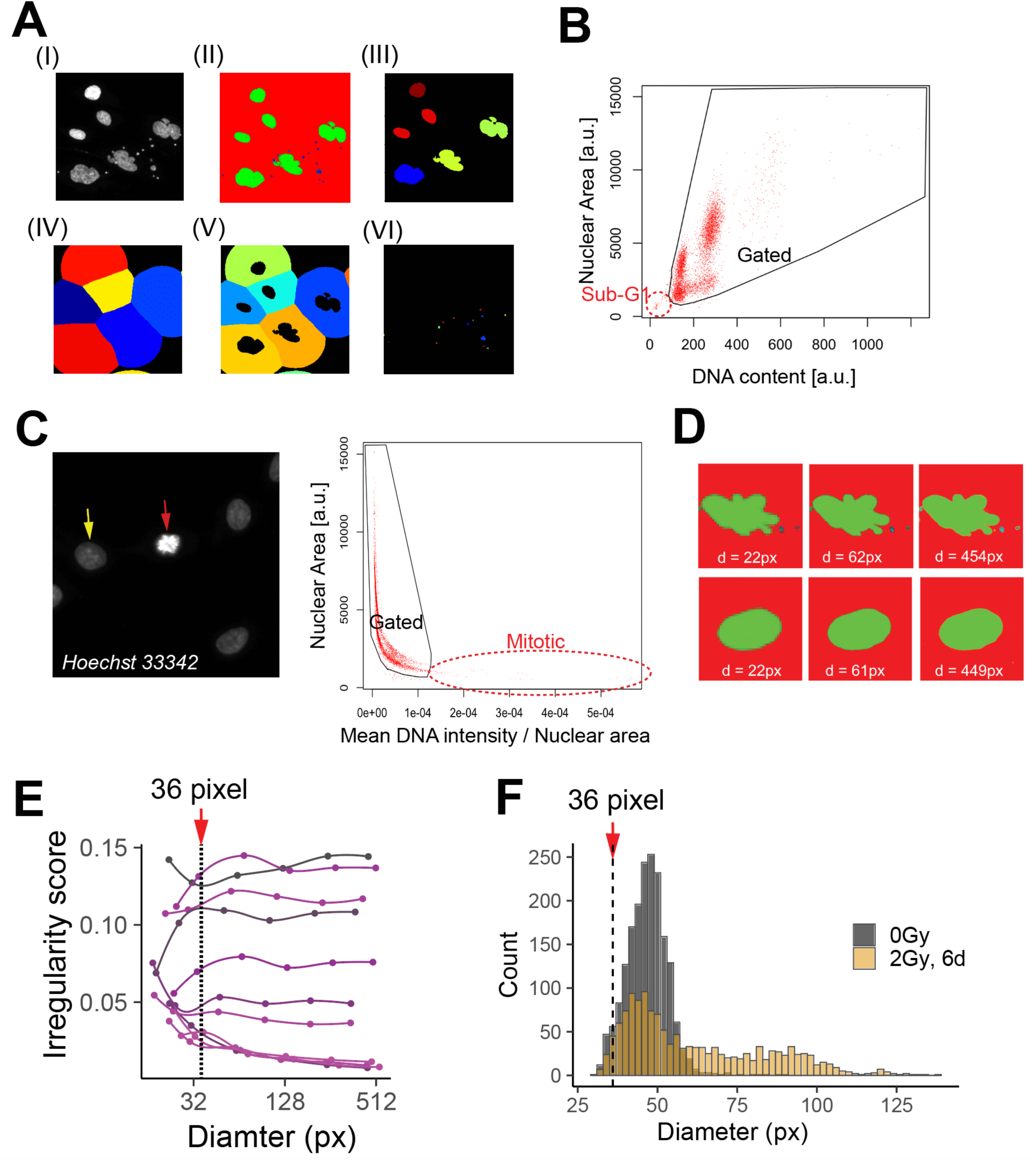
Strategies for nuclear segmentation, data cleaning, and quantification of irregularity score. **(A)** Example of image segmentation process. (I) Max-projected Hoechst 33342 staining. (II) Probability map output by ilastik. Red, background; green, primary nucleus; blue, micronuclei. (III) Segmented primary nuclei. (IV, V) Defining peripheral region for each primary nucleus. Each nucleus was expanded by certain pixels (IV), and primary nucleus was masked from each expanded area to define peripheral region for each primary nucleus (V). (VI) Micronuclei (if any) within nuclear peripheral regions were segmented by processing the probability map (II). **(B)** Gating example of objects based on DNA content and nuclear area. Objects with sub-G1 DNA content, demarcated by the red dashed circle, were excluded from the gating. **(C)** Gating example to exclude mitotic cells. Left: DNA staining in a mitotic cell (red arrow) and interphase cells (yellow arrow to one of them). Right: objects with exceptionally high mean DNA intensity divided by nuclear area, demarcated by the red dashed ellipse, were left out from the gating. **(D)** Examples of two nuclei with different pixel resolutions. d, nuclear diameter in pixels (EquivalentDiameter in CellProfiler). px, pixel. **(E)** Examples of how pixel resolution affects quantified irregularity scores. Irregularity scores from the same nucleus were connected by a line. **(F)** Example histogram of nuclear diameter in cells under non-damaged and damaged conditions.

**Figure S8:**
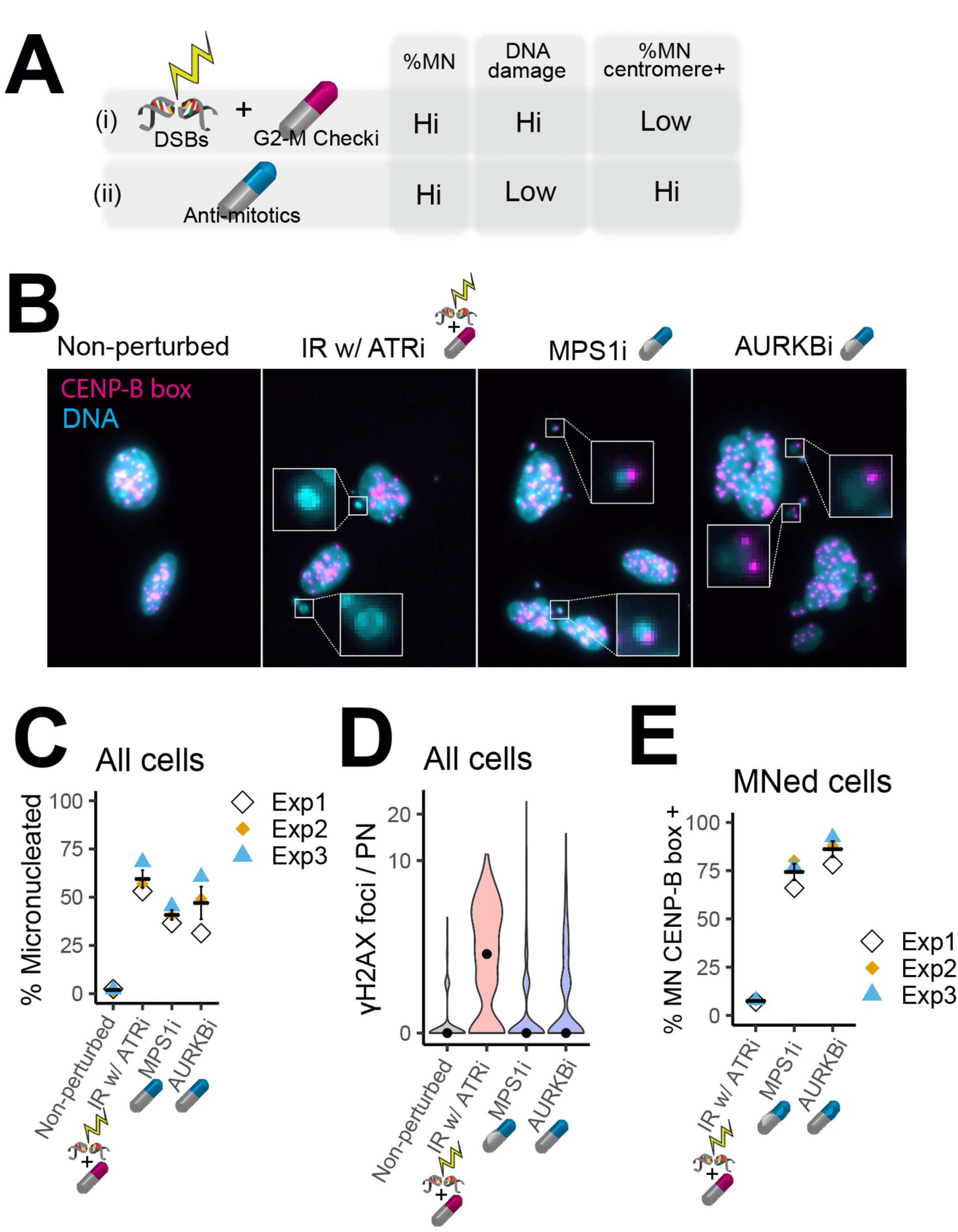
Centromere presence in micronuclei informs origin of mitotic error. **(A)** Expected effect of forced mitosis under DSBs (i) or antimitotics with chromosome missegregation properties (ii) on micronucleation, DNA damage, and micronuclear centromere (MN centromere). **(B)** Representative centromere staining in micronucleated cells generated under the indicated treatments. Cells were treated for one day before centromere quantification using PNA FISH assay targeting the repetitive centromeric sequence CENP-B box. **(C)** Percentage of micronucleation in cells treated as in B (n = 3 biological replicates with >350 cells each). Error bar, SEM. **(D)** Number of γ-H2AX foci per primary nucleus in cells treated under the indicated conditions (n = 3 biological replicates with >350 cells each) – data from C. **(E)** Percentage of micronucleated cells with micronuclear CENP-B box signal, in cells treated under the indicated conditions (n = 3 biological replicates with >140 micronucleated cells each) – data from C. Error bar, SEM.

